# A cellular taxonomy of the bone marrow stroma in homeostasis and leukemia demonstrates cancer-crosstalk with stroma to impair normal tissue function

**DOI:** 10.1101/556845

**Authors:** Baryawno Ninib, Przybylski Dariusz, Monika S. Kowalczyk, Kfoury Youmna, Severe Nicolas, Gustafsson Karin, Mercier Francois, Tabaka Marcin, Hofree Matan, Dionne Danielle, Papazian Ani, Lee Dongjun, Rozenblatt-Rosen Orit, Regev Aviv, David T Scadden

**Affiliations:** Center for Regenerative Medicine, Massachusetts General Hospital, Boston, MA 02114, USA; Harvard Stem Cell Institute, Cambridge, MA 02138, USA; Department of Stem Cell and Regenerative Biology, Harvard University, Cambridge, MA 02138, USA; Klarman Cell Observatory, Broad Institute of Harvard and MIT, Cambridge, MA 02142, USA; Department of Biology and Koch Institute, MIT, Boston, MA 02142, USA; Department of Convergence Medical Science, Pusan National University School of Medicine, Yangsan 50612, Republic of Korea; Howard Hughes Medical Institute, Chevy Chase, MD 20815, USA

**Author notes:** Correspondence (AR), (D.T.S.). These authors contributed equally (Baryawno Ninib, Przybylski Dariusz, Monika S. Kowalczyk).

## Abstract

Stroma is a poorly defined non-parenchymal component of virtually every organ with key roles in organ development, homeostasis and repair. Studies of the bone marrow stroma have defined individual populations in the stem cell niche regulating hematopoietic regeneration and capable of initiating leukemia. Here, we use single-cell RNA-seq to define a cellular taxonomy of the mouse bone marrow stroma and its perturbation by malignancy. We identified seventeen stromal subsets expressing distinct hematopoietic regulatory genes, spanning new fibroblastic, and osteoblastic subpopulations. Emerging acute myeloid leukemia resulted in impaired osteogenic differentiation and reduced production of hematopoietic regulatory molecules necessary for normal hematopoiesis. Thus, cancer can affect tissue stroma in which they reside to disadvantage normal parenchymal cells. Our taxonomy of the regulatory stromal compartment provides experimental support for a model where malignant clone is not a destroyer of normal tissue but an architect of it, remodeling tissue stroma to enable emergent cancer.

## INTRODUCTION

The tissue microenvironment of stem cell niches maintains and regulates stem cell function through cellular interactions and secreted factors (1, 2). Hematopoiesis provides a paradigm for understanding mammalian stem cells and their niches, with pivotal understanding from numerous *in vivo* studies on the critical role of several non-hematopoietic niche cells as regulators of hematopoietic stem cell (HSC) function (3-7).

One major component are multipotent mesenchymal stem/stromal cells (MSCs), non-hematopoietic cells derived from the mesoderm with potential to differentiate into bone, fat and cartilage *in vitro* (8). While MSCs are found in most tissues, their diversity and lineage relationships are incompletely understood. For instance, several subtypes of MSCs have been described in bone specialized niches that regulate HSC maintenance. Most of these cells are located in the perivascular space and associated with either arteriole or sinusoidal blood vessels, produce key niche factors such as Cxcl12 and Stem Cell Factor (SCF, also known as *Kitl*) (9), and are identified by Leptin receptor [Lepr-cre] (4, 10), Nestin [Nes-GFP] (6) or Ng2 (*Cspg4*) [NG2-CreER] (5) expression. However, it remains unclear if these markers delineate distinct or overlapping cell populations.

Other non-hematopoietic cells, including endothelial cells (ECs) and MSC-descendent osteolineage cells (OLCs), also play roles as niche cells. Endothelial cells produce Cxcl12, SCF, and other niche factors and are critical regulators of HSC function (4, 11-16). OLCs are critical for HSC homing after lethal irradiation and bone marrow transplantation (17), modulate hematopoietic progenitor function and lineage maturation (10, 18, 19), and dysfunction in some of them has been implicated in myelodysplasia and leukemia development (20-23).

However, despite extensive studies, the HSC niche remains incompletely defined in terms of its cellular and molecular composition, limiting our ability to prospectively isolate and functionally characterize niche cells. Previous profiling studies of MSCs were performed in bulk and relied on reporter genes to purify cell populations (9), which may either analyze a mixed population (if marker expression is more promiscuous than assumed), only cover a subset (if the marker is overly specific), or fail to detect unknown or transient states.

Here, we use single cell RNA-seq (scRNA-seq) to chart a comprehensive census of the non-hematopoietic cells of the bone marrow. We identified 17 distinct subsets and their associated gene signatures, key transcription factors (TFs), cell surface markers and secreted factors for each of the major cell types, and defined stromal cells expressing key niche factors at steady-state hematopoiesis. We further inferred their differentiation relationships, and characterized diversity within specific cell types. Finally, we dissect the influence of cancer development in the bone marrow microenvironment at the global level and shed light on cellular and molecular abnormalities as acute myeloid leukemia develops.

## RESULTS

### A scRNA-seq census of bone marrow stroma identifies MSCs, OLCs, chondrocytes, fibroblasts, ECs and pericytes, and their differentiation relations

To explore the cellular composition of the stroma of the mouse bone marrow, we profiled non-hematopoietic bone marrow cells by scRNA-seq. We enriched stroma cells from six C57Bl/6 mice at the age of 8-10 weeks by FACS (**Fig. 1a**), through selective depletion of hematopoietic cells and immune lineage cells as labeled by CD45, lineage and Ter119 (**Supplementary fig. 1a**), profiled them by droplet-based scRNA-seq (**Methods**), and retained 30,543 high quality profiles for further analysis (**Supplementary fig. 1b**, **Methods**). We partitioned the cells into 33 clusters by unsupervised clustering (**Supplementary fig. 1c**, **Methods**), which we annotated *post-hoc* (**Methods**), and distinguished subsets spanning 20,896 non-hematopoietic cells from 9,647 cells in hematopoietic clusters (**Fig. 1b**, **Supplementary fig. 1c,d**). Each of the non-hematopoietic clusters contained cells from all six mice (**Supplementary fig. 1e**).

**Fig 1.**
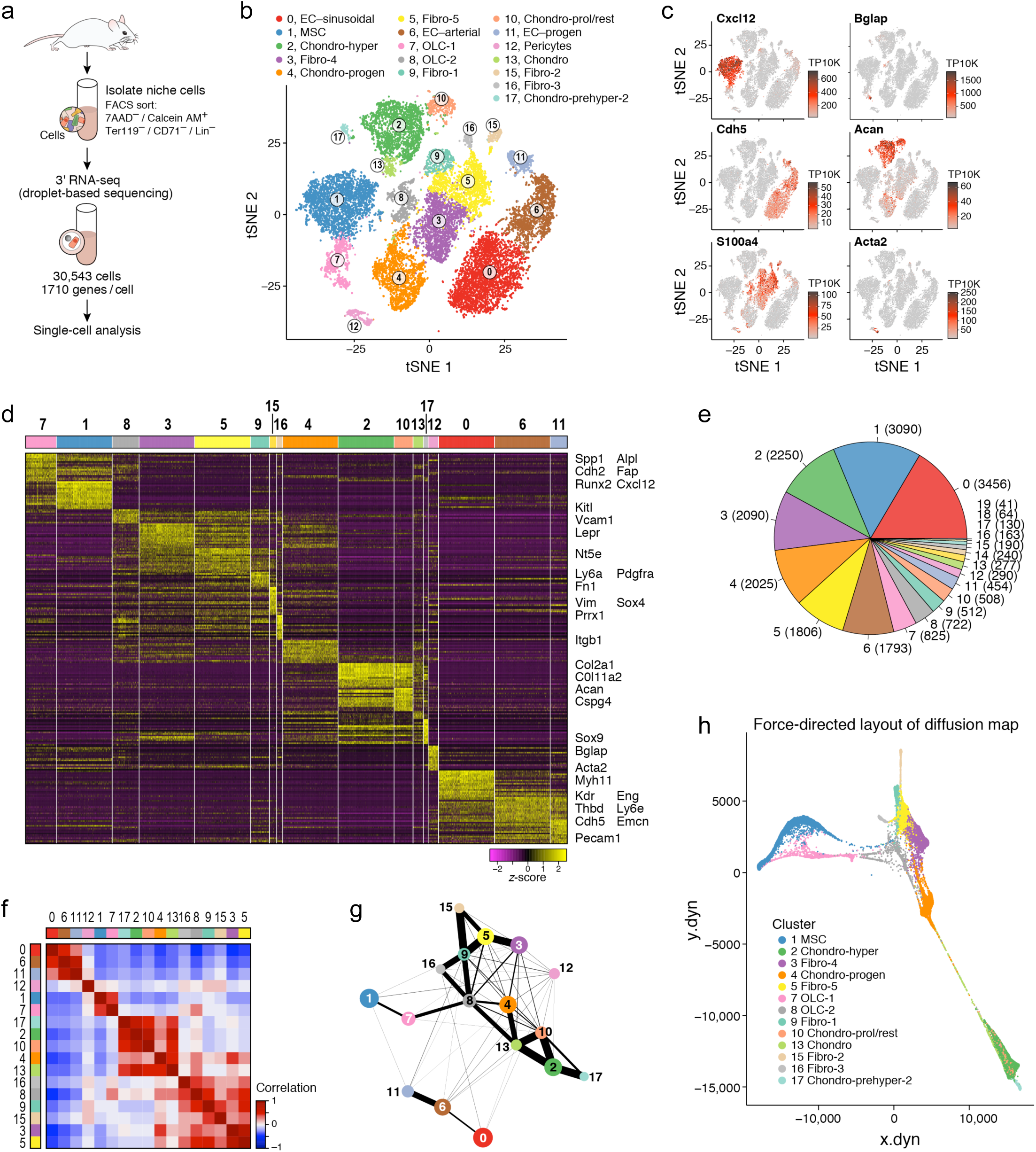
A single cell atlas of the mouse stroma. (**a**) Study Overview. (**b**,**c**) Seventeen BM stroma cell clusters. t-SNE of 20,551 non-hematopoietic cells colored by clustering and annotated *post hoc* (b) or by expression (color bar, TP10K) of key marker genes genes for each six major cell types (c). (**d**) Cluster signature genes. Expression (row-wide z-score of ln of TP10K) of top differentially expressed genes (rows) across the cells (columns) in each cluster (color bar, top, as in B). The cells from the largest clusters were down-sampled for visualization. Key genes are highlighted on right. (**e**) Number of cells in each subset. Color bar as in B. (**f-g**) Differentiation relations between cells or clusters. (**f**) Pearson’s correlation (color bar) between the average gene expression profile of each pair of clusters (rows, coloumns, color code as in b). (**g**) Relatedness (edges; width indicates strength) between clusters (nodes, colored as in B) based on cluster graph abstraction (AGA; **Methods**). (**h**) A force directed layout embedding (FLE) of the cells (dots) from a diffusion map (50 components) computed with the cells from strongly connected clusters (as indicated in g and f, without clusters 1, 6, 11, 12).

Clustering of only the non-hematopoietic cells partitioned them into 17 clusters (**Fig. 1b**), spanning MSCs, OLCs, chondrocytes, fibroblasts, ECs and pericytes and possible transitional states (**Fig. 1c-h**, **Supplementary fig. 1f-h**). In particular, there were: (**1**) MSCs (cluster 1, expressing *Lepr* (10) and *Cxcl12* (24)); (**2**) two OLC subsets (Cluster 7 and 8, expressing osteocalcin (*Bglap*) (25)); (**3**) four chondrocyte subsets (clusters 2, 10, 13 and 17, annotated by expression of the chondrocyte markers, Aggrecan (*Acan)* (26) and *Col2a1* (27)); (**4**) five fibroblasts subsets (Cluster 3, 5, 9, 15 and 16; expressing specific protein-1 (*S100a4*) (28), collagens and extracellular matrix (ECM) proteins); (**5**) three EC subsets (Clusters 0, 6, 11; annotated by expression of the pan-endothelial marker Ve-cadherin (*Cdh5*) (29)); (**6**) one pericyte cluster (Cluster 12, expressing *Acta2* (30)); and (**7**) likely differentiating cells (Clusters 4 and 8, expressing markers of chondrocytes, osteoblasts and fibroblasts), indicating a potential differentiation pathway emanating from MSCs with distinct origin from other osteoblastic cells. Across subsets, chondrocytes, fibroblasts and OLCs had the highest proliferation status as estimated by expression of a cell cycle signature (**Supplementary fig. 1i**, **Methods**).

To help resolve differentiation relations between the cells we used correlation of average cluster profiles (**Fig. 1f**), graph abstraction (31) (**Fig. 1g**), and diffusion map analysis (32) (**Fig. 1h**, **Supplementary fig. 1f**). We discuss these below in the context of each lineage and differentiation path.

We validated two of the most abundant subsets, MSCs and three EC subsets by FACS, showing that the four clusters can be partitioned prospectively by combining antibodies that labels ECs (CD31, Sca-1, CD34) or MSCs (CD106/*Vcam1*) (**Supplementary fig. 1g,j,k**) in combination with the antibodies we used previously to isolate stroma from immune (Lin-) and erythroid (Er-) cells (**Supplementary fig. 1a**).

### MSCs producing HSC regulators form four subsets that span a differentiation continuum

The prevailing model of MSCs in the bone marrow is that of a multipotent stem cell that can differentiate into bone cells, adipocytes and chondrocytes (33), but their exact identity remains unclear. In particular, while many protein markers have been proposed and deployed in mice (*e.g.*, Cxcl12, LepR, Nes, NG2 (*Cspg4*) (4-6, 10, 34), or human (CD73 (*Nt5e*), CD106 (*Vcam1*) (35), CD105 (*Eng*) (33), CD90 (*Thy1*) (36)), there is no single accepted combination and some may also be expressed by pericytes, ECs and chondrocytes (8). We hypothesized that more comprehensive transcriptional profiles will better identify subsets and relate them to legacy markers.

We annotated cluster 1 (**Fig. 2a**) as LepR^+^ MSCs/CAR cells with preadipocytic features (4, 24, 34, 37). First, the cells highly expressed *Lepr*, a perivascular MSC marker (4, 9), adiponectin (*Adipoq*), a gene highly expressed in preadipocytes (38) (**Fig. 2c**, **Supplementary fig. 2c**), and *Cxcl12, Kitl* and *Angpt1* (**Fig. 2b,c**, **Supplementary fig. 2c**), key niche factors produced by bone marrow niche cells. Notably, *Cxcl12, Kitl* and *Angpt1* were also expressed albeit to a lower level in a subset of OLCs (cluster 7, **Fig. 2a,b**), which were contiguous with cluster 1 in the inferred differentiation maps (**Fig. 1g,h**), suggesting continuous transitions in niche functions on the MSC-OLC differentiation path. The cells expressed additional MSC markers (*Nt5e, Vcam1* and *Eng*) as well as *Grem1*, a BMP antagonist, a marker of mouse skeletal stem cells with bone, cartilage and reticular marrow stromal cell potential (39) (**Supplementary fig. 2a,c**). Other previously proposed MSC markers, such as *Thy1, Ly6a* (Sca-1) and the perivascular MSC markers NG2 (*Cspg4*) and Nestin (*Nes*) (5, 6), were not expressed in cluster 1, but in other, non-MSC cells (**Supplementary fig. 2a,b**).

**Fig. 2.**
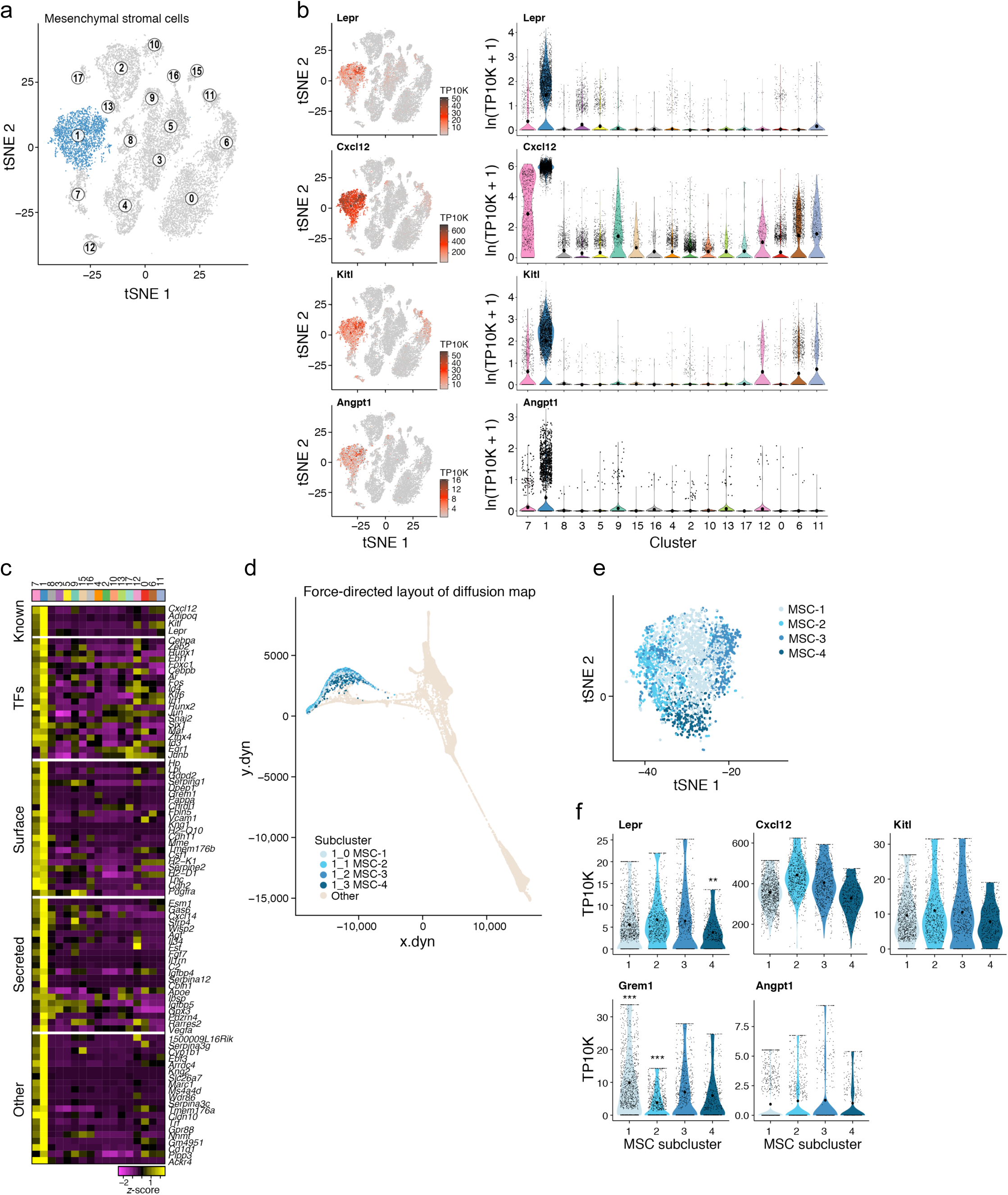
Four subsets of HSC regulator-producing MSCs form a differentiation continuum. (**a-c**) Signature genes for MSC in the stroma atlas. (a,b) tSNE of **Figure 1b** colored by cluster-1 (MSC) membership (a), or by expression (color bar, TP10K) of key MSC marker genes (b, left) (right), along with the distributions of expression levels (ln(TP10K+1), *y* axis) for the same genes across the 17 clusters of **Figure 1b** (*x* axis). (**c**) Expression (row-wide z-score of ln of average TP10K) of top differentially expressed genes (rows) in the cells of each cluster (columns) (color bar, top, as in **Figure 1b**), ordered by five gene categories (labels on left). (**d**-**f**) MSC subclusters span a differentiation continuum. (d,e) Continuous transition across the subclusters. Four MSC subclusters color coded on the FLE of the diffusion map in **Figure 1h** (e) or on a zoom-in to the MSC cluster in **Figure 2a**. (f) Distributions of expression levels (TP4K, *y* axis, censored scale) for select marker genes across the 4 sub-clusters.

Within cluster 1 MSCs, we further distinguished four major subsets (MSC-1 to 4) by sub-clustering and diffusion trajectory analysis (40) (**Fig. 2d,e**, **Supplementary fig. 2d**,**e Methods**). While *Lepr, Kitl* and *Angpt1* were comparably expressed in all four subsets (albeit lowest in MSC-4), they were distinguished by the expression of *Cxcl12, Grem1* and multiple other genes (**Fig. 2f**, **Supplementary fig. 2e**). In particular, one subset (MSC-4) had high expression of osteolineage cell-specification genes (*Runx2, Sp7* (osterix), *Col1a1* and *Alpl* (41, 42)) (**Supplementary fig. 2e,f**), suggesting differentiation towards the osteoblastic lineage, supported by the continuous transition inferred between cells from the MSC and OLC-1 clusters in differentiation trajectories (**Fig. 1g,h**). Indeed, OLCs that include osteoprogenitors and mature osteoblasts, can be derived from MSCs (8).

### Inferred lineage relations suggest two osteolineage subsets of different differentiation origins and distinct hematopoietic support potential

To further analyze osteolineage differentiation, we focused on cells from the MSC-4 subsets, along with cluster 7 (OLC-1) and 8 (OLC-2) cells (**Fig. 3a**) that express the osteoblast maturation marker *Bglap* (**Fig. 3b**, **Supplementary fig. 3a**) (25), and form a continuum in the diffusion map (**Fig. 1h)**. All three clusters differentially express *Runx2*, the master regulator TF controlling the commitment of MSCs to OLCs (41), and both OLC-1 and OLC-2 further express *Sp7*, its downstream target TF, which together are required for both early and late differentiation (**Fig. 3b,c**, **Supplementary fig. 3a**). Furthermore, *Bglap* is expressed in a subset of the clusters’ cells (**Fig. 3b**), along with other osteoblastic maturation makers (*Alpl, Col1a1, Spp1* and *Pth1r* (41)) (**Fig. 3b,c**, **Supplementary fig. 3b**).

**Fig. 3.**
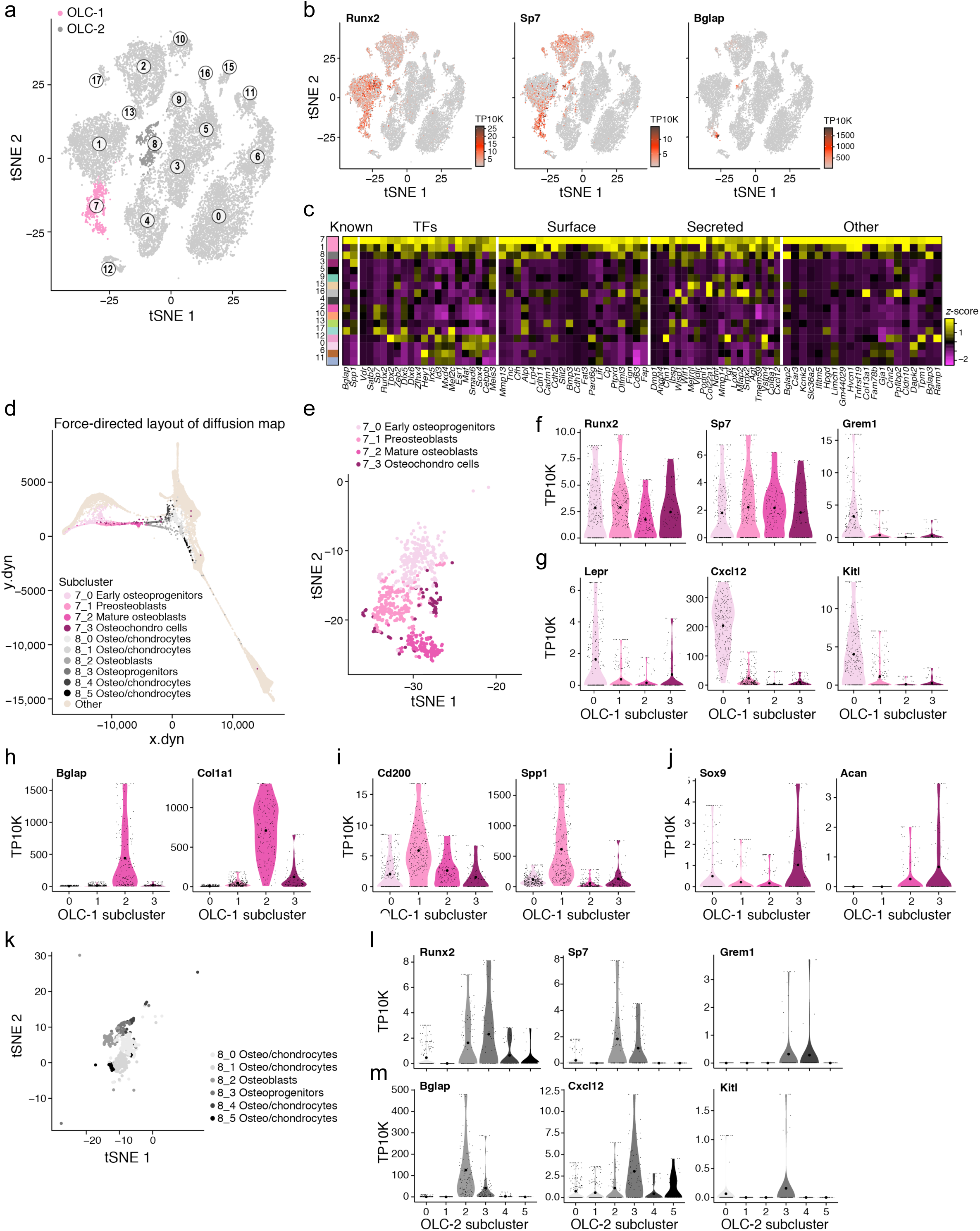
Two osteolineage subsets of distinct differentiation origins and hematopoietic support potential. (**a-c**) Two OLC subsets. (a,b) tSNE of **Figure 1b** colored by cluster 7 (OLC-1) and cluster 8 (OLC-2) membership (a), or by expression (color bar, TP10K) of key MSC marker genes (b). (c) Expression (column-wide z-score of ln of average TP10K) of top differentially expressed genes (rows) ordered by four gene categories (labels on top) in the cells of each cluster (columns, color bar, left, as in **Figure 1b**). (**d**) Continuous transition across the subclusters. OLC-1 and OLC-2 subclusters color coded on the FLE of the diffusion map in **Figure 1h** (d). (**e-j**) OLC-1 subsets. (e) A zoom-in to the OLC-1 cluster in **Figure 2a** labeled by four subclusters and annotated *post hoc* (legend). (f-j) Distributions of expression levels (TP10K, *y* axis, censored scale) for select marker genes across the 4 OLC-1 sub-clusters. (**k-m**) OLC-2 subsets. (k) A zoom-in to the OLC-2 cluster in **Figure 2a** labeled by six subclusters and annotated *post hoc* (legend). (l,m) Distributions of expression levels (TP10K, *y* axis, censored scale) for select marker genes across the 6 OLC-2 sub-clusters.

Whereas MSC-4 reflects committed osteolineage MSCs, the cells in OLC-1 span a continuum of osteoblastic differentiation as indicated by diffusion map (**Fig. 3d,e**) and differential gene expression (**Supplementary fig. 3c**) analyses, including, in order: (**1**) osteolineage committed early progenitors that were in a cell transition-state from MSC-4 (**Fig. 2e and** **3d**,**e**), expressing *Runx2, Sp7, Grem1, Lepr, Cxcl12* and *Kitl* (**Fig. 3f,g**, **Supplementary fig. 3c**), but not osteoblast maturation markers (*Bglap* and *Col1a1*; **Fig. 3h**); (**2**) late osteoprogenitor cells (*e.g.*, preosteoblasts), highly expressing the skeletal stem cell and osteoprogenitor marker CD200 (43, 44) and *Spp1* (**Fig. 3i**), a marker of later stages of osteoblastic differentiation (41), but lowly expressing osteoblast maturation markers (**Fig. 3h**); and (**3**) mature osteoblasts, expressing *Bglap* and *Col1a1* (**Fig. 3h**). In addition, the subcluster included (**4**) chondrocytes, which are likely transitioning into osteoblastic cells, through endochondral ossification (45, 46). These expressed low levels of *Sox9*, a TF essential for chondrocyte differentiation (47, 48), and *Acan*, a chondrocyte marker gene (26) (**Fig. 3J**).

Remarkably, the OLC-2 cells highlighted a distinct subset of OLCs from those found in a continuum from MSCs. Along the continuum spanned by OLC-2 cells (**Fig. 3d,k and Supplementary fig. 3d**), we observed: (**1**) Osteoprogenitors (8_3), expressing *Runx2, Sp7*, and *Mmp13*, the matrix metalloproteinase that regulates the calcification and degradation of cartilage during the endochondral ossification process (49) (**Fig. 3l and Supplementary fig. 3e**), but low levels of *Bglap* and *Sox9*, (**Fig. 3m**, **Supplementary fig. 3e**); (**2**) Osteoblasts (8_2), expressing OLC markers (*Runx2*, Sp7, *Spp1, Bglap* and *Pth1r*, **Fig. 3l**,**m and Supplementary fig. 3e**); and (**3**) osteo/chondrocytes (8_0, 8_1, 8_4 and 8_5) that were in the process of being ossified into bone (45, 46). Notably, only OLC-1 cells express key hematopoietic regulatory cytokines (albeit 8_3 in OLC-2 express relatively low levels *Cxcl12*) (**Fig. 3m**), suggesting that there are two osteolineage subsets of distinct differentiation origins and with distinct hematopoietic support potential.

### Characterization of chondrocyte cell differentiation

We next examined the differentiation of chondrocytes. Cartilage is formed by chondrocytes that are derived from condensed MSCs that differentiate into chondrocyte progenitors. Cartilage development occurs at the growth plate, where chondrocytes transition between proliferating and resting states, differentiate to hypertrophic chondrocytes, which in turn terminally lose their capacity to proliferate, and finally are ossified into bone (50).

Cells in five clusters (Clusters 2, 4, 10, 13, 17) all expressed the chondrocyte lineage genes *Sox9, Acan* and *Col2a1* (**Fig. 4a,b**), formed a sequence in the overall diffusion map (**Fig. 4c**) and distinguished: (**1**) Proliferating and resting chondrocytes (Cluster 10), expressing *Col2a1* (51) (**Fig. 4a,b**, **Supplementary fig. 1i**); (**2**) prehypertrophic chondrocytes that started the maturation process (Clusters 2 and 17), co-expressing *Ihh, Pth1r* (52), *Mef2c* (53) (**Fig. 4a**, **Supplementary fig. 4a-c**), (**3**) hypertrophic chondrocytes (Cluster 2), co-expressing *Runx2, Ihh, Mef2c* and *Col10a1* (**Fig. 4a**, **Supplementary fig. 4a-c**) (53); as well as (**4**) chondrocyte progenitor cells that are in transition into cluster 8 (Cluster 4) (**Fig. 4c**, **1h**), expressing *Grem1* (**Supplementary fig. 4b**), OLC markers (*Runx2, Sp7, Alpl, Spp1*) (**Fig. 3b**, **Supplementary fig. 3b**), and chondrocyte markers (*Sox9, Acan*; **Fig. 4b**).

**Fig. 4.**
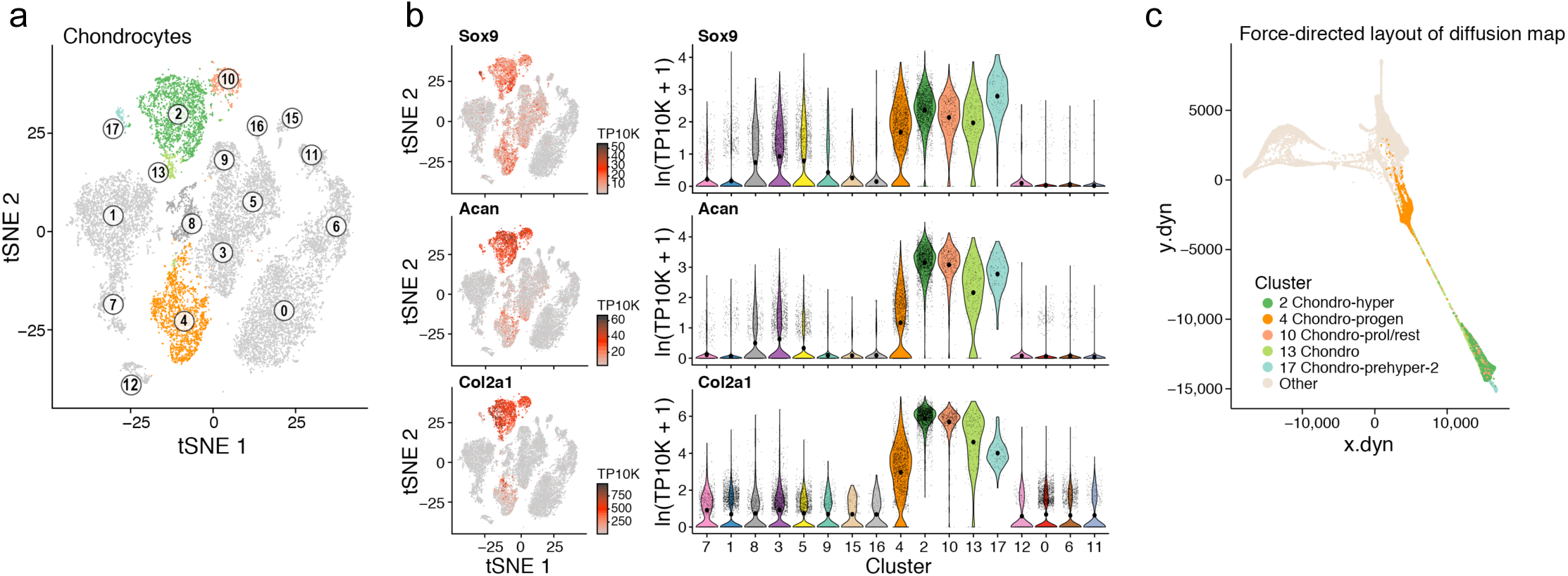
Chondrocyte subsets highlight differentiation paths. (**a-c**) Subsets across chondrocyte differentiation. (a,b) Signature genes for chondroid subsets. tSNE of **Figure 1b** colored by membership in five chondroid clusters (a), or by expression (color bar, TP10K) of key marker genes (b, left) (right), along with the distributions of expression levels (ln(TP10K+1), *y* axis) for the same genes across the 17 clusters of **Figure 1b** (*x* axis). (c) Continuous transition across the chondroid subclusters. The five subclusters color coded on the FLE of the diffusion map in **Figure 1h**.

### A bone marrow derived fibroblast subset expresses hematopoietic niche regulators

Fibroblasts are cells of mesenchymal origin that are ubiquitous in the bone marrow, and consist of phenotypically and functionally distinct subpopulations. Currently, fibroblasts are predominantly identified by *Fn1, Pdgfra*, Fibroblast Specific Protein-1 (*S100a4*) and *Acta2* expression (54). Due to the paucity of distinctive markers and similar morphology and phenotypes, they are commonly confounded in the bone marrow with MSCs and perivascular cells, limiting the accuracy of functional studies (55). We hypothesized that we can better define the precise identity and function of bone marrow fibroblasts by their transcriptional profiles.

Five fibroblasts clusters (3, 5, 9, 15, 16, **Fig. 5a**) all highly expressed the fibroblast specific genes Fibronectin-1 (*Fn1)*, Fibroblast Specific Protein-1 (*S100a4*), Decorin (*Dcn*), and Semaphorin-3C (*Sema3c*) (**Fig. 5a,b**), lowly expressed the chondrocyte genes *Sox9, Acan*, and *Col2a1* (**Fig. 4b**), and were distinguishable from both MSCs and pericytes. Cells in distinct clusters expressed different collagens, indicating specialized roles in connective tissue formation (56) (*Col1a1* and *Col1a2* in clusters 3, 5 and 9; *Col22a1* in clusters 15 and 16; **Supplementary fig. 5a**).

**Fig. 5.**
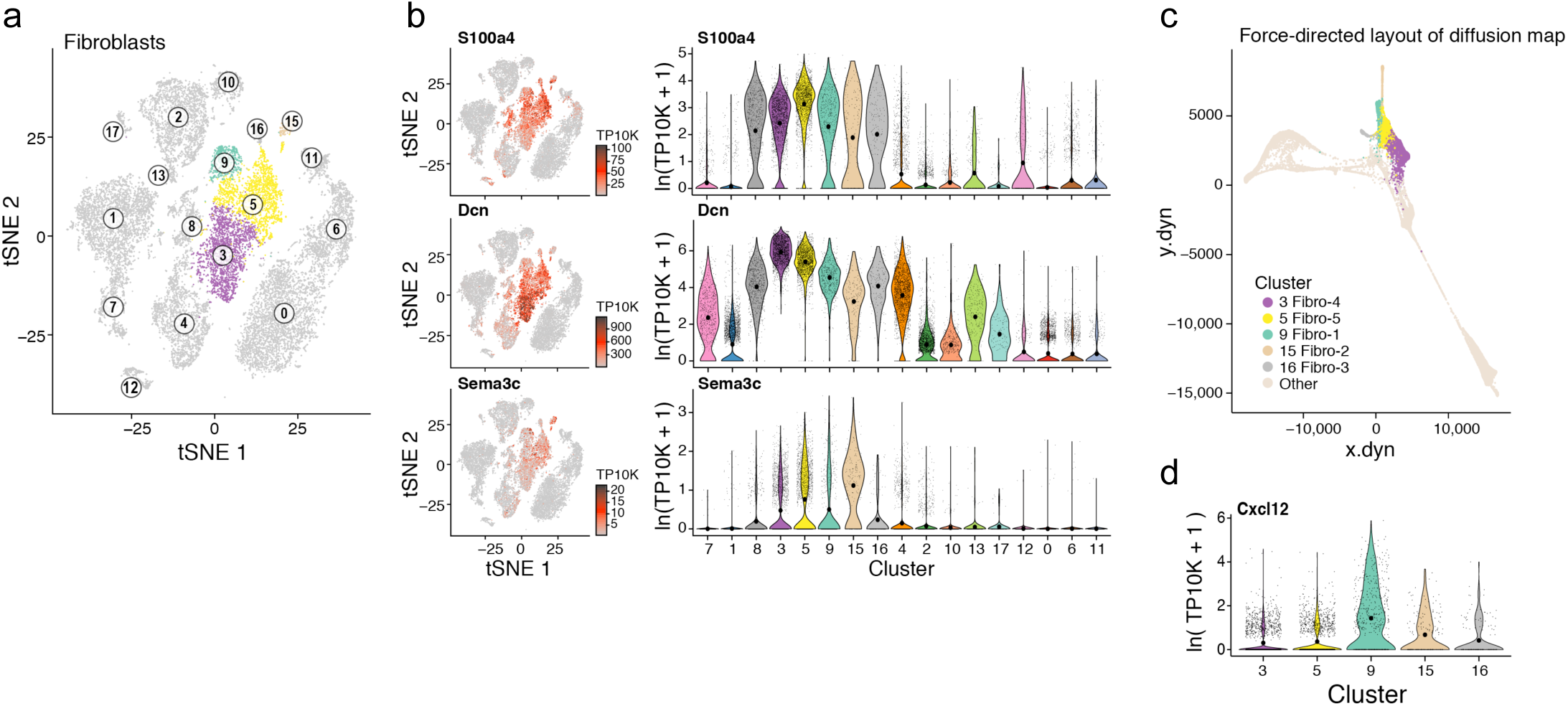
Fibroblasts subsets highlight hematopoiesis support. (**a-c**) Fibroblast subsets. (a,b) Signature genes for fibroblast subsets. tSNE of **Figure 1a** colored by membership in five fibroblast clusters (a), or by expression (color bar, TP10K) of key marker genes (b, left) (right), along with the distributions of expression levels (ln(TP10K+1), *y* axis) for the same genes across the 17 clusters of **Figure 1b** (*x* axis). (c) Relations between the fibroblast subclusters. The five subclusters color coded on the FLE of the diffusion map in **Figure 1h**. (**d**) Some fibroblast clusters that express the key niche factor *Cxcl12.* Distribution of *Cxcl12* expression level (ln(TP10K+1), *y* axis) across the 5 fibroblast clusters (*x* axis).

**Fibroblasts-1** (cluster 9) and **2s** (cluster 16) had MSC characteristics and some may provide niche regulatory functions. Their cells expressed the progenitor marker *Cd34* (**Supplementary fig. 1g**) and MSC markers (*Ly6a, Pdgfra, Thy1* and *Cd44*; **Supplementary fig. 4a,b**), but not EC or pericytes genes (*Cdh5, Acta2* (**Fig. 1c)**), suggesting these fibroblasts are MSC-like. Fibroblast-1s also express the hematopoietic regulatory cytokines *Cxcl12* and *Angpt1* (**Fig. 2b**, **4d**), indicating a possible niche regulatory function that resembles CAFs that facilitate cancer growth, metastasis (57), and immunosuppression through Cxcl12 (58).

**Fibroblasts-3**, **4**, **and 5s** were related to the tendon/ligamentous junction, from tenocyte progenitors to tendon/ligament cells. Fibroblast-3 (cluster 16), Fibroblast-4 (Cluster 3) and Fibroblast-5 (cluster 5) co-expressed *Sox9* (**Fig. 4b**) and the TF Scleraxis (*Scx*) (**Supplementary fig. 4c**), which regulate differentiation in tenocytes and ligamentocytes in tendons and ligaments (59). Fibroblast-4 and -5s further expressed key bone and cartilage genes, including the osteogenic *Spp1* (**Supplementary fig. 3b**) and the chondrocyte genes *Cspg4, CD73* (*Nt5e*) (**Supplementary fig. 2a**) and Cartilage Intermediate Layer Protein (*Cilp*) (**Supplementary fig. 4c**). The three clusters formed a continuum from Fibroblasts-3s to Fibroblast-4s and -5s (**Fig. 4c**). Thus, fibroblast-3s may be tenocyte progenitors (59), whereas Fibroblast-4s and -5s consist of tendon/ligament cells that are located close to the chondro-tendinous/ligamentous junction (59).

### EC progenitor express higher levels of niche factors compared to sinusoidal and arteriolar vascular ECs

Bone marrow derived endothelial cells (BMECs) are a heterogeneous cell population (60) from either arteriolar or sinusoidal blood vessels, where they act as critical regulators of HSC function through secretion of niche factors (4, 11-13, 15, 61-63).

We identified three BMEC subsets (Clusters 11, 6, and 0) – annotated as EC progenitors, arterial BMECs, and sinusoidal BMECs, respectively. All subsets expressed *Pecam1, Cdh5, Kdr*, and *Emcn* (64) (**Fig. 6a-c**), and related along a continuum (**Fig. 6d**). In particular, *Cd34*, commonly expressed in primitive endothelial and hematopoietic cells (65, 66), was highly expressed in both cluster 6 and cluster 11, indicating an immature cell state. Diffusion map analysis (40) (**Fig. 6d**) suggests that cluster 11 cells are EC progenitors that differentiate into cluster 6 cells. We annotated cluster 6 and 0 cells as arterial and sinusoidal BMECs (aBMECs and sBMECs), respectively, by marker expression: Cluster 0 cells had high expression of *Flt4* (Vegfr-3) and low expression of *Ly6a* (Sca-1), and cluster 6 cells had the opposite pattern (**Supplementary fig. 6a**). This was consistent with previous reports that sinusoidal BMECs are Vegfr-3^+^/Sca-1^-^, while arterioles that branch from arterial BMECs are Vegfr-3^-^/Sca-1^+^ (13, 14).

**Fig. 6.**
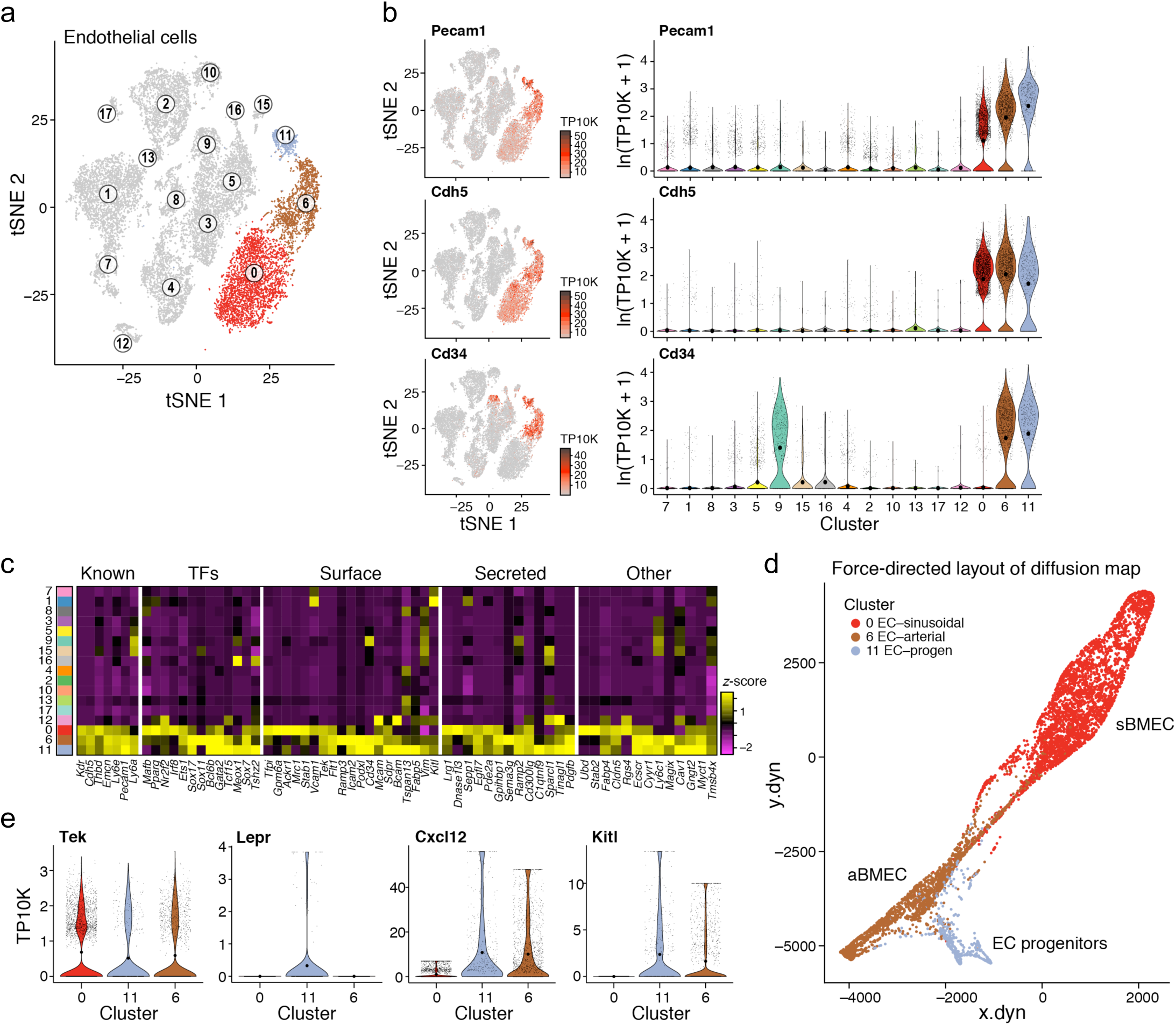
EC progenitors express higher levels of niche factors. (**a-c**) Three EC subsets. (a,b) tSNE of **Figure 1b** colored by membership in three EC clusters (a), or by expression (color bar, TP10K) of key marker genes (b, right), along with the distributions of expression levels (ln(TP10K+1), *y* axis) for the same genes across the 17 clusters of **Figure 1b** (*x* axis) (b, left). (c) Expression (column-wide z-score of ln of average TP10K) of top differentially expressed genes (columns) ordered by five gene categories (labels on top) in the cells of each cluster (rows, color bar, left, as in **Figure 1b**). (**d**) Continuous transition across the subclusters. FLE of a diffusion map of ECs, color coded by cluster membership. (**e**) Key marker genes. Distributions of expression levels (ln(TP10K+1), *y* axis) for selected genes across the three clusters (*x* axis).

The three subsets expressed different ligands and secreted factors. All three expressed the endothelial tyrosine kinase receptor, Tie2 (*Tek* gene product) for angiopoietin ligands. However, the hematopoietic factors, *Kitl* and *Cxcl12* were most abundant in the EC progenitors, aBMECs expressed *Cxcl12* with minimal *Kitl*, and sBMECs did not express either factor (**Fig. 6e**, **Supplementary fig. 6c**). Thus, EC progenitors may be a distinctively relevant subpopulation within ECs, that regulates hematopoietic stem and progenitor cell function (4, 67).

### NG2^+^ and Nestin^+^ pericytes are distinct from LepR^+^ MSCs

NG2^+^ and Nestin^+^ pericytes have been proposed as critical regulators of HSC function through production of *Cxcl12* and *Kitl* (5, 6). Notably, under homeostasis, HSCs reside in perisinusoidal or periarteriolar locations predominantly found in close proximity to LepR-cre positive cells, also shown to produce *Cxcl12* and *Kitl* (9, 68). However, determining whether NG2^+^ and Nestin^+^ pericytes share developmental origins and functional properties with perivascular LepR^+^ MSCs (30) has been challenging as there is currently no single marker that identifies them without overlapping with MSCs (30).

We annotated a distinct cluster of pericytes (Cluster 12), by the co-expression of the classical markers Nestin (*Nes*), NG2 (*Cspg4*), α-smooth muscle actin (*Acta2*), myosin (*Myh11*) and *Mcam* (30) (**Fig. 7a-c**, **Supplementary fig. 7a**). The pericytes differentially express *Jag1*, the Notch signaling ligand, and *Il-6*, both implicated in promoting HSC maintenance (3, 69) (**Supplementary fig. 7c**). Notably, cells in this pericyte cluster express only very low levels of *Lepr*, in contrast to the high expression of *Lepr* in MSCs (**Fig. 2b,c**). Thus, perivascular MSCs and pericytes are clearly distinguishable.

**Fig. 7.**
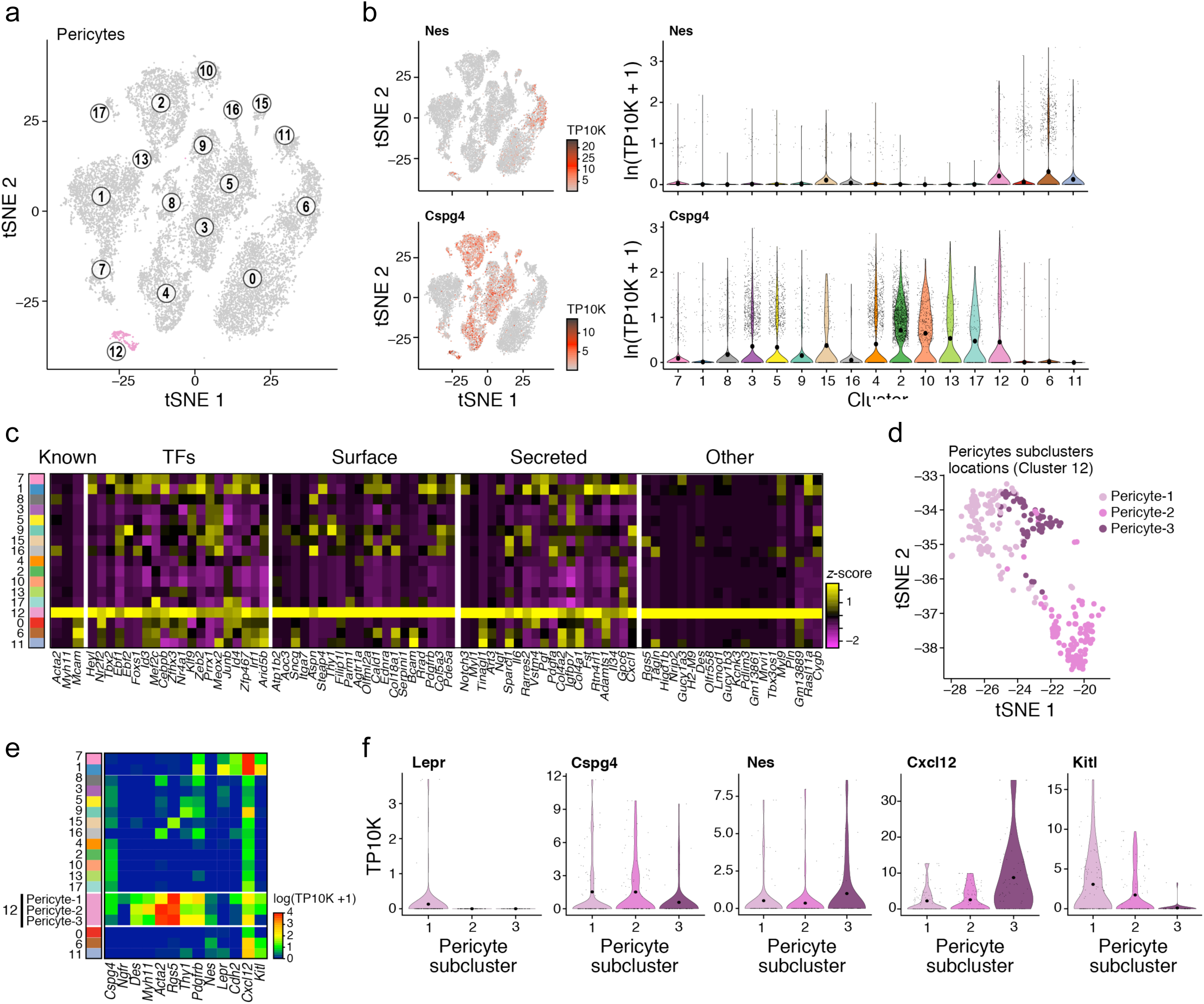
Three distinct subpopulations of pericytes, distinguished from LepR^+^ MSCs. (**a-c**) Signature genes for Pericytes in the stroma atlas. (a,b) tSNE of **Figure 1b** colored by cluster-12 (pericyte) membership (a), or by expression (color bar, TP10K) of key pericyte marker genes (b, left) (right), along with the distributions of expression levels (ln(TP10K+1), *y* axis) for the same genes across the 17 clusters of **Figure 1b** (*x* axis). (c) Expression (column-wide z-score of ln of average TP10K) of top differentially expressed genes (columns) in the cells of each cluster (rows) (color bar, left, as in **Figure 1b**), ordered by five gene categories (labels on left). (**d**) Pericyte subclusters color coded on the zoom-in of to the pericyte cluster in **Figure 7a**. (**e**) Average expression (ln(TP10K+1)) of select hematopoietic niche genes (x axis) in 17 clusters of **Figure 1b** (y axis). Three pericyte sub-clusters indicated. (**f**) Distributions of expression levels (TP10K, *y* axis, censored scale) for select marker genes across the 3 sub-clusters.

Within the pericytes, we identified three subsets (**Fig. 7d** and **Supplementary fig. 7b**). Pericyte-1 cells, distinguished by relatively higher expression of *Cdh2* and specific expression of MSC surface marker *Ngfr* (CD271) (**Fig. 7e**), expressed *Lepr, Cxcl12, Cspg4* and *Nes* (albeit at low levels for the four genes) and *Kitl*, and may reflect a subset of LepR^+^ cells in the pericyte pool that contribute to HSC regulation (**Fig. 7e,f**). Conversely, LepR^-^ pericytes express either *Cspg4* (NG2, Pericyte-2s) or *Nes* (Pericyte-3s), which have previously characterized, respectively, periarteriolar cells (NG2-CreER/Nestin-GFP^high^ (5, 70)) and perisinusoidal cells (Nestin-GFP^+^ (6)), co-localizing with HSC and serving as HSC niche cells. In marked contrast to MSCs, none of these subsets expresses high levels of both *Cxcl12 and Kitl* (**Fig. 7e,f** and **Fig. 2f**). Therefore, while LepR^+^ cells, *Cspg4*^*+*^ (NG2) or *Nes*^*+*^ cells have been defined as HSC niche cells, they are molecularly distinct. *Cspg4* (NG2) and *Nes* expressing pericytes are not among the broad range of LepR^+^ cells (4, 10), and their distinct expression of *Cxcl12* and *Kitl* suggests they may have distinct niche functions.

### Systemic survey of changes in bone marrow stromal cells induced by primary MLL-AF9 leukemia

Several bone marrow stromal cell populations that regulate hematopoiesis, can, when perturbed, lead to niche-initiated myelodysplasia or leukemia in animal models (20, 21, 23, 71). Moreover, myeloid malignancies can remodel their niche to support malignant growth (72-74).

To comprehensively assess changes in the stroma during malignant growth, we used a mouse model (75), where we engrafted primary MLL-AF9 knock-in bone marrow donor cells (4-6 week old mice) into congenic wild-type recipient mice where leukemia evolved, and compared them to comparably conditioned bone marrow transplants from mice without MLL-AF9 (**Methods**). More than 6 months of normal hematopoiesis elapsed in both populations before leukemia was detected in the MLL-AF9 bearing mice. Mice showing signs of emerging leukemia, enlarged spleens (**Supplementary fig. 8a**) and leukemic blasts by FACS scatter and Cd11b^+^/Gr1^-^ expression within the myeloid compartment in peripheral blood (**Supplementary fig. 8b-c**) were sacrificed ∼6-10 months post-transplantation.

We profiled by scRNA-seq 12,456 bone marrow stromal cells from MLL-AF9 (MLL) mice (n=4) and 10,548 cells from matched control transplanted mice (n=5). We clustered the cells from all mice together, and assigned cell-type identity based on gene signatures from our steady-state analysis (**Fig. 8a,b**, **Supplementary fig. 8d**). We then tested for statistically significant differences in the proportions of each cluster/subset.

**Fig. 8.**
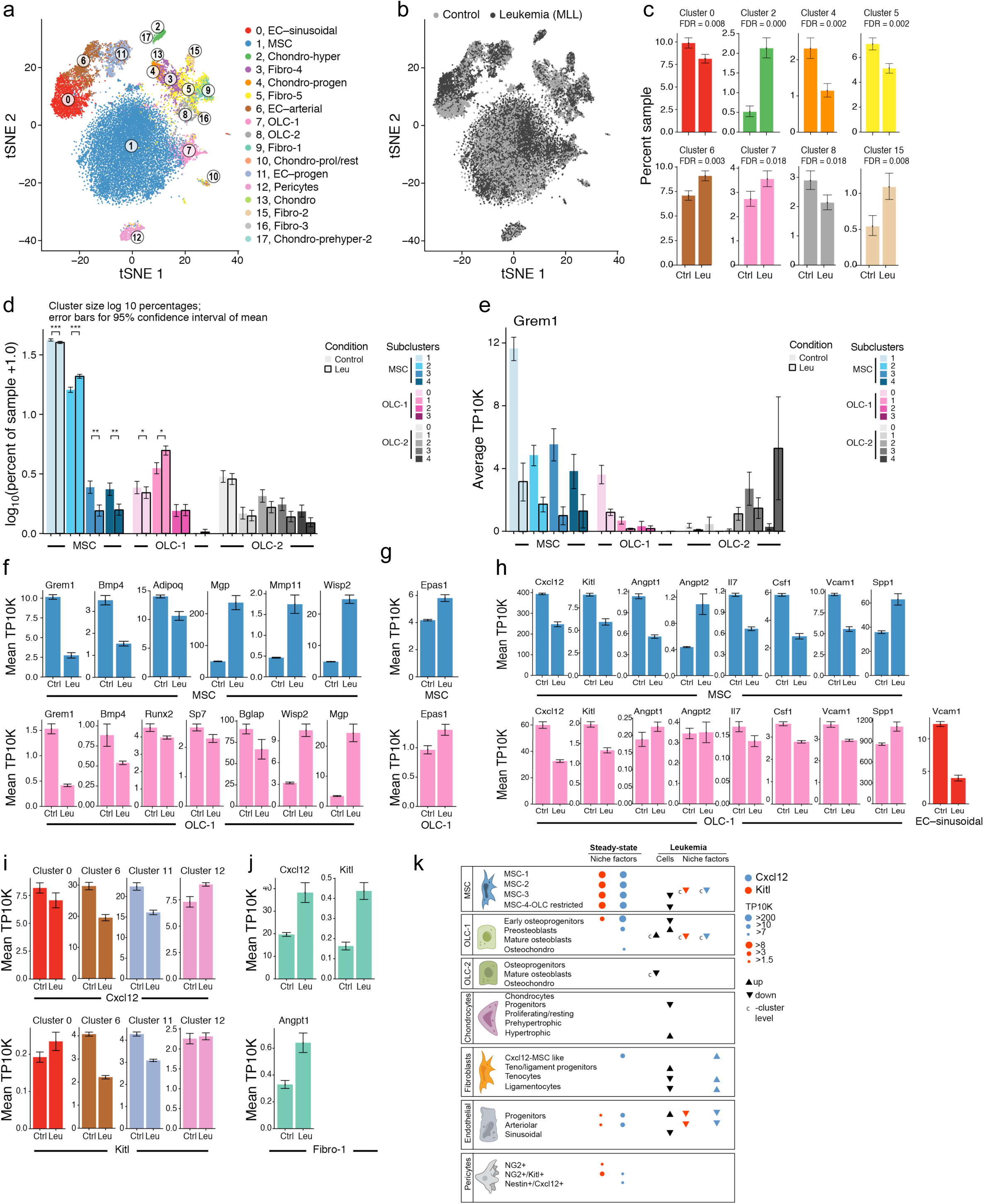
Remodeling of the bone marrow stroma in leukemia. (**a**,**b**) A census of the leukemic bone marrow stroma. tSNE of 23,004 cells (dots) from control and leukemic bone marrow colored by cluster assignment (as in Figure 1b), or by condition (control: light grey; leukemic: dark grey). (**c**,**d**) Compositional changes in BM stroma. Binomial fit mean as percent of cells (y axis) assigned to specific cluster (c) or to MLC and OLC subclusters (d) among control or leukemic samples (*x* axis). Error bars: 95% confidence interval of the binomial fit mean. (**e**) Changes in Grem1 expression in MSCs and OLCs. Average of samples (TP10K, y axis) in MSC and OLC sub-clusters (x axis). Error bars: SEM. (**f-h**) Changes in BM niche remodeling, hypoxia and hematopoietic regulator genes in MSCs and OLC-1s in leukemia. Average expression (TP10K, y axis) of BM niche modelling genes in MSCs (top) and OLC-1s (bottom). Error bars: SEM. (**i**,**j**) Changes in *Cxcl12, Kitl* and *Angpt1* expression in sBMEC, aBMEC, progenitor ECs, pericytes or MSC-like fibroblasts. Average expression (TP10K, y axis) of the denoted genes in specific cell clusters. Error bars: SEM. (**k**) A census of the bone marrow stroma in homeostasis and leukemia. Horizontal boxes: broad cell types, with subsets noted. Circles: expression levels of key niche factors, *Kitl* and *Cxcl12*. Dark triangles: changes in subset proportion in Leukemia. Colored triangles: changes in relative expression in leukemia. ‘*C’* -mark next to triangles indicates a cell type level change (*i.e.*, refers to all subsets).

### Remodeling of OLCs, ECs, chondrocytes, and fibroblasts proportions in leukemia

We detected significant changes in the proportions of key subsets in leukemia. These often reflected coupled effects in OLCs (increase in OLC-1, reduction in OLC-2), ECs (reduction in sBMECs, increase in aBMECs), chondrocytes (increase in cluster 2, decrease in 4) and fibroblasts (decrease in Fibroblast-3s, increase in Fibroblast-4s) (**Fig. 8c**, **Supplementary fig. 8e**).

The shift in OLCs, including reduction in more mature bone cells, while maintaining Nes^+^ pericytes, is consistent with previous studies of leukemia, where OLC lineage dysfunction and loss of mature cells is caused by leukemia cells and favorable for their growth (76-79). Moreover, proportions also varied within the OLC-1 subset, with a significant increase in preosteoblasts (subcluster 7-1) but not in other cells (**Fig. 8d**). While the overall proportion of MSCs was unchanged (**Supplementary fig. 8e**), the relative proportions *within* MSCs were impacted by leukemia, with significant increase in MSC-2 and a concomitant reduction in MSC-3 and MSC-4 (committed osteolineage MSCs) (**Fig. 8d**).

Overall, the concomitant increase in osteoprogenitors (*e.g.*, preosteoblasts) and decrease in committed osteolineage MSCs suggests that leukemia induced a block in osteolineage development (80). The changes observed in OLCs and MSCs, as well as ECs, are consistent with the reported abnormalities seen in AML patients, where leukemia induces vasculature remodeling that is accompanied by reduced osteocalcin (*Bglap*) serum levels, growth deficiency, and impaired osteogenesis (76, 80-82).

### Leukemia compromises osteogenic and adipogenic differentiation pathways in MSCs and OLCs

Analyzing changes in intrinsic gene programs within each cluster (**Table S1**, **Supplementary fig. 8f-i**) highlighted blunted adipogenic and osteogenic differentiation programs in MSCs and OLCs. Cells in both MSC and OLC clusters downregulated the skeletal stem cell marker *Grem1* (39), across all MSC subclusters and in OLC progenitors (**Fig. 8e**), as well as the BMP pathway, including *Bmp4* (GSEA (83) (GSEA qval=0.027, **Table S2**) (84), and the early osteolineage cell-specification gene *Sp7* (**Fig. 8f)**. Mature osteoblasts within OLCs further downregulated *Runx2*, an early marker of osteogenesis, and *Bglap1, Bglap2* and *Bglap3*, late markers of osteogenesis (85) (**Fig. 8f**, **Supplementary fig. 8f**). MSC cells further downregulated *Adipoq* (**Fig. 8f**, **Supplementary fig. 78**) (86) and other genes in the white adipocyte differentiation (GSEA qval=0.029, **Table S2**), and the PPAR signaling pathways (GSEA qval=0.022, **Table S2**) (87). This is consistent with reports that a compromised adipocytic niche induced by AML leads to imbalanced stem and progenitor regulation and not favorable for normal blood cell production (88). Furthermore, *Wisp2*, which has been implicated in restricting MSCs to an undifferentiated state (89, 90) was upregulated in all MSC subsets and OLC progenitors (within OLCs) (**Supplementary fig. 78**,**j)**, further supporting compromised differentiation of MSCs and OLCs. Conversely, key genes that inhibit bone formation and calcification were induced in MSCs and OLCs (and others), including *Mgp* (**Fig. 8f**) (inhibits calcification, (91)); *Igbfp5* and *Igfbp3* (negative regulators on bone formation (92, 93), **Supplementary fig. 8g**), and genes in the ECM degradation pathway (GSEA qval=0.04, **Table S2**), including *Mmp2, Mmp11* and *Mmp13* (**Supplementary fig. 8g**). Taken together, these data are consistent with AML inducing alterations that affect bone formation and breakdown.

The cell profiles further support a model (94) where hypoxia contributes to the undifferentiated state of the MSCs and OLCs. Hypoxia pathway genes (GSEA qval=0.004, **Table S2**) were induced in MSCs and OLCs, including the key regulator Hif-2a (*Epas1*) (**Fig. 8g** and **Supplementary fig. 8h**), but not *Hif-1a* (**Supplementary fig. 8j**), consistent with reports that suppression of bone formation and osteoblast activity is directly linked to Hif-2a and not Hif-1a (95). Notably, the *Wisp2* promoter is regulated by Hif-2a (96).

### Leukemia broadly impairs normal hematopoiesis-regulatory expression

The changes in the stroma may contribute to altered support of normal blood cell growth, through deregulation of the expression of key HSC niche factors, especially *Cxcl12* and *Kitl* across MSCs, aBMECs, the earliest OLCs (subcluster 7-0) and EC progenitors (**Fig. 8h,i**); conversely, *Cxcl12, Kitl* and *Angpt1* were upregulated in Fibroblast-1s (**Fig. 8j**). These findings are consistent with prior observations of AML reducing HSC persistence and localization to the bone marrow (97), and that Cxcl12-secreting-CAFs (similar to Fibroblast-1s) are associated with a cancer promoting phenotype in breast cancer (57, 58).

Among other factors, *Spp1*, a negative regulator of HSC pool size (98) and HSC proliferation (99), which is correlated with poor prognosis in AML patients (100), was upregulated in MSCs, OLCs, ECs, fibroblasts and pericytes (**Fig. 8h**, **Supplementary fig. 8f**). MSCs down regulated other HSC niche factors including *Angpt1*, an agonist of *Tek* receptor expressed on ECs and HSCs (101) (and upregulated its antagonist *Angpt2* (102)); factors that promote lymphoid (*Il7*) or myeloid (*Csf1*) differentiation (103, 104) (105) and *Vcam1*, a regulator of HSC homing to bone marrow (106) (**Fig. 8h**). Thus, the osteogenic differentiation block induced by leukemia in MSCs and OLCs, was further accompanied by a loss of HSC niche factor production by multiple cell types.

## DISCUSSION

While it is now well appreciated that non-parenchymal cells in the stroma of a tissue play key physiological roles, in forming niche cells (3, 7), the full repertoire of cells that comprise stroma has remained elusive in all complex adult tissues, including the bone marrow. Here, we systematically characterize the non-hematopoietic cells of the mouse bone marrow into six broad cell types with 17 cell subsets (**Fig. 8k**), with discrete distinctions, differentiation continuums and HSC niche regulatory function. The five cell types are mesenchymal stromal cells (MSC), osteolineage cells, chondrocytes, endothelial cells, and pericytes. Unexpected subtypes of cells were found within these cell groups such as two subsets of osteoblasts, four chondrocyte populations and three subsets of endothelial cells. Distinct profiles of hematopoietic regulatory gene expression further distinguished subsets of cells and indicated likely participation in hematopoietic regulation, often disrupted by the emergence of leukemia (**Fig. 8k**).

These findings provide a clear set of definitions and tools, including refined understanding of the relation between markers and cell types (and their limitations), clarifies prior controversies, and leads to several unexpected observations. Here we highlight some of these key findings.

First, osteoblasts segregated into two subsets that appear to arise from distinct lineage trajectories. Only one of these, which emerges from MSCs, expresses HSC regulatory genes and does so at the osteoprogenitor stage of development.

Second, one of the five fibroblasts subsets we identified, termed Fibroblasts-1, expresses *Cxcl12*. Cxcl12-expressing fibroblasts have been implicated in aggressive solid tumors (57, 58). We thus hypothesize that they may participate in aggressive bone metastases; future studies can test this possibility.

Third, ECs were the most abundant cell type in bone marrow stroma, and include a distinctive, immature subset that is enriched for expression of hematopoietic regulators. This is in marked contrast to the large sinusoidal endothelium subset that minimally expresses the HSC niche factors, Cxcl12 and Kitl, despite prior reports that peri-sinusoidal positioning of HSC is common (68).

Fourth, we clarified that LepR^+^ cells are extremely common across multiple cell types, including osteolineage, endothelial, pericyte and fibroblastic cells. Thus, the use of LepR-driven Cre to alter particular genes could impact their expression across many cell types, and must be interpreted with extreme care, especially where LepR-Cre is expressed throughout development, as commonly done to date (107). We further showed that LepR expressing cells are quite distinct from Nestin or Cspg4 (Ng2) producing cells. The latter have been previously reported as important HSC niche cells, suggesting that subtle functional distinctions may be discerned between HSCs associated with the distinct niches these cells may represent. Within MSCs, where *LepR* expression is abundant, *Grem1* is disproportionately expressed in particular MSC subsets. *Grem1*^*+*^ cells have been described as functionally different than *LepR+* cells in their relative ability to make specific mature marrow subsets such as adipocytes (39, 107). As the expression of *Grem1* does not track precisely with *LepR*, these may indeed represent cell populations with graded functional capabilities.

Fifth, pericytes were inferred as of distinct lineage from the MSC populations reported to be perivascular (6) and there was discordant expression of *Cxcl12* and *Kitl* among pericyte subpopulations. Pericytes with abundant *Cxcl12* also express *Cspg4* and *Nes*, but have little detectable *Kitl* or *LepR*. Future studies can assess if these reflect functional distinctions in hematopoietic support, and how this sub-population tracks with periarteriolar, quiescent HSCs (5).

Finally, the presence of acute myeloid leukemia distorted the stromal compartment in select and specific ways. Osteogenic differentiation blockade occurs in MSCs and OLCs, is associated with a hypoxia signature, and accompanied by a cell intrinsic bone remodeling phenotype and disturbed production of hematopoietic regulatory factors that affect normal hematopoiesis (*Cxcl12, Kitl, Angpt1* and *Spp1*). While most studies show that osteoblast numbers are either reduced (77, 79, 80) or increased (73), depending on type of leukemia model used, our study supports a loss of bone maturation phenotype and function. A parenchymal tumor affecting the maturation of tissue stromal cells does suggest a distinct type of cross-interaction between emerging cancer cells and their mesenchymal neighbors. It may be that differentiation blockade is not restricted in a cell autonomous manner to cancer cells, but may extend to the broader cell context of a cancerous tissue. Additional cell intrinsic changes associated with AML in specific subclusters impacted HSC support factors expressed by MSCs, osteoprogenitors, endothelial progenitors and arteriolar ECs, with marked reduction in expression of genes important for HSC retention and persistence (*Cxcl12, Kitl* and *Angpt1*), hematopoietic maturation (*Il7* and *Csf1*) and cell adhesion (*Vcam1*).

These findings are consistent with a model where emerging cancer cells influence the stromal cells in the tissue they inhabit. They can alter differentiation patterns of those stromal cells, changing the complexity of cell types that are thought to play critical roles in governing tissue homeostasis. Furthermore, the malignant cells in our study reduced the expression of regulatory signaling molecules known to be essential for normal hematopoietic function. In so doing, the malignant cells create a microenvironment no longer as conducive to normal hematopoietic cell production and, thereby, impairing the parenchymal cells with which they compete. These data provide experimental support for a paradigm in which the establishment of a malignant clone within a tissue shapes the features of the stromal landscape of that tissue. The result compromises stromal cell support of normal parenchymal cells, fundamentally altering the competitive landscape of the tissue to disadvantage normal cells. In this way, cancer cells act not as fully independent and destructive rogues, but rather as self-serving architects of their neighborhood pushing out normal occupants by creating a less supportive environment.

Our stroma cell census will now allow a clearer and more consistent definition of the ways in which specific stromal cells contribute to homeostasis and aberrant hematopoiesis, and provide a foundation for developing stromal-targeted therapies in hematologic disease. It also offers experimental evidence for tumor evolution in which cross-communication between parenchymal and stromal elements may influence the emergence of cancer.

## ACKNOWLEDGMENTS

N.B. was funded by the Swedish Research Council and The Childhood Cancer Foundation in Sweden. M.S.K. was supported by Charles A. King Trust Postdoctoral Research Fellowship Program and the Simeon J. Fortin Charitable Foundation. L. Gaffney assisted with Fig. preparation. Work was supported by the Klarman Cell Observatory and HHMI (AR), NIH DK107784 and the Gerald and Darlene Jordan Professorship (DTS). We thank professor Henry Kronenberg for helpful comments.D.T.S. is a director and shareholder of Magenta Therapeutics, Agios Pharmaceuticals, Clear Creek Bio, Red Oak Medicines and LifeVaultBio; he is a shareholder of Fate Therapeutics. A.R. is a founder of Celsius Therapeutics and a member of the SAB for ThermoFisher Scientific, Driver Group and Syros Pharmaceuticals. M.S.K. is currently employed by Celsius Therapeutics.

## Author contributions

N.B., D.P., M.K., A.R., and D.T.S. conceived and designed the project. N.B. carried out bone marrow stromal cell harvesting, FACS sorting and FACS analysis. N.S.,K.G. and D.L. contributed with harvesting of bone marrow stroma and FACS sorting.M.K. performed scRNA-seq, with help from D.D., and guidance from O.R.R. D.P. performed computational analyses, with help and code contributions from M.H. and M.T. N.B., D.P., M.K., Y.K., N.S. and D.T.S contributed to data interpretation. N.B., Y.K. and F.M. performed bone marrow transplants, breeding and generation of leukemic mice. A.P. performed bleeding and peripheral blood analysis. N.B., D.P., M.K., A.R. and D.T.S. wrote, reviewed and edited the manuscript. A.R. and D.T.S. supervised the project. All authors commented on the manuscript.

## METHODS

### Mice

The MLL-AF9 knock-in mice (75) and CD45.1 (STEM) mice (108) were described previously. Littermates were used as controls for all experiments involving MLL-AF9 knockin and CD45.1 (STEM) mice. Male C57BL/6 mice (CD45.2, Jackson Laboratory) at age 6-8 weeks were employed as transplant recipients and for steady-state scRNA-seq experiments. All animal experiments were performed in accordance with national and institutional guidelines. Mice were housed in the Massachusetts General Hospital (MGH) Animal Research Facility on a 12 hour light/dark cycle with stable temperature (22° C) and humidity (60%). All procedures were approved by MGH Internal Animal Care and Use Committee.

### Bone marrow transplantation and generation of leukemic mice

To generate leukemic mice, we first crossed the MLL-AF9 knock-in mice (75) with the CD45.1 (STEM) mice (109) to generate donor chimeric CD45.1.2 mice. Mice positive for the MLL-AF9 fusion transgene were used as donors, and littermates negative for the MLL-AF9 fusion transgene were used as controls. Mice were sacrificed via CO2 asphyxia; tibiae and femurs were harvested and excess soft tissue was eliminated. Bones were crushed and washed in PBS and passed through a 70μm filter into a collection tube and1x10^6^ whole bone-marrow cells were transplanted by retro-orbital injection into 6-8 weeks old male CD45.2 C57BL/6 recipient mice. One day prior to transplantation, mice were subjected to whole body irradiation (2 x 6Gy) with a 6-hour interval from a ^137^Cs source. Monthly retro-orbital bleeding was performed on isoflurane anesthetized mice and blood was withdrawn using heparinized capillaries and collected into EDTA containing tubes to prevent coagulation. Complete blood counts were done using the Element Ht5 Auto Hematology analyzer. Subsequently, RBCs were lysed as described previously and cells were stained in PBS, 2% FBS using the following antibodies: CD45.2-APCCy7 (Biolegend, Ref#109824, clone 102), CD45.1-FITC (Biolegend, Ref# 110706, clone A20), CD11b-Alexa Fluor 700 (Ref# 101222, clone M1/70), GR1-Brilliant violet 570 (Biolegend, Ref#108431, clone RB6-8C5), B220-eFluor450 (eBioscience, Ref#48-0452-82, clone RA3-6B2), and CD3e-APC (eBioscience, Ref#17-0031-83, clone 145-2c11), in addition to 7-Aminoactinomycin D (7AAD; ThermoFisher Scientific, Ref#A1310) for viability to monitor donor chimerism within the different lineages and the appearance of leukemic blasts characterized by a distinct scatter and lower GR1 and CD11b expression within the myeloid compartment. Leukemic mice were determined by the combination of disease symptoms, white blood cell counts and appearance of leukemic blasts.

### Isolation of bone marrow stroma cells

To obtain bone marrow niche cells for scRNA-seq, mice were sacrificed via CO2 asphyxia. Bones (femur and tibia) were harvested and placed in Media 199 (ThermoFisher Scientific, Ref#12350039) supplemented with 2% Fetal Bovine Serum (FBS, ThermoFisher Scientific, Ref#10082147). Muscle and tendon tissue was removed and bone marrow was flushed. Niche cells from the bone marrow fraction were isolated by digestion with 1 mg/mL STEMxyme1 (Worthington, Ref#LS004106) and 1 mg/mL Dispase 1 (ThermoFisher Scientific, Ref#17105041), in Media 199 supplemented with 2% FBS for 25 min at 37°C. Niche cells from the bone fraction were isolated by gently crushing and cutting bones into small fragments and digested in the same digestion mix as the bone marrow for 25 min, at 37°C with agitation (120 rpm). After digestions, both fractions were filtered through a 70μm filter into a collection tube (Fisher Scientific, Ref#08-771-2), pooled into one sample, and erythrocytes lysed in ACK-lysis buffer (ThermoFisher Scientific, Ref#C1430) for 5 minutes on ice. Cells were then stained in Media 199 supplemented with 2% FBS for FACS cell sorting.

### FACS enrichment of bone marrow niche cells

For flow cytometry and FACS, cells were resuspended in Media 199 supplemented with 2% FBS and stained for Ter119-APC (eBioscience, Ref#17-5921-82, clone TER-119), CD71-PECy7 (Biolegend, Ref#113812, clone RI7217), CD45-PE (eBioscience, Ref#12-0451-82, clone 30-F11), CD3-PE (Biolegend, Ref#100206, clone 17A2), B220-PE (Biolegend, Ref#103208, clone RA3-6B2), CD19-PE (Biolegend, Ref#115508, clone 6D5), Gr-1-PE (Biolegend, Ref#108408, clone RB6-8C5), and Cd11b-PE (Biolegend, Ref#101208, clone M1/70) for 30 minutes on ice. Dead cells and debris were excluded by FSC, SSC, DAPI (4’,6-diamino-2-phenylindole, Life Technologies, Ref#D3571) and Calcein AM staining profiles and Calcein AM (Life Technologies, Ref#C1430). FACS and cytometry was performed on a BD FACSAria II sorter, and sorted bone marrow niche cells were collected in Media 199 supplemented with 2% FBS and 0.4% UltraPure BSA (Life Technologies, Ref#AM2616). Bone marrow stroma was enriched by sorting of live cells (7-AAD-/Calcein+) negative for erythroid (CD71/Ter119) and immune lineage markers (CD45/CD3/B220/CD19/Gr-1/CD11b).

### Single cell RNA-seq

Single cells were encapsulated into emulsion droplets using Chromium Controller (10x Genomics). scRNA-seq libraries were constructed using Chromium Single Cell 3’ v2 Reagent Kit according to the manufacturer’s protocol. Briefly, post sorting sample volume was decreased and cells were examined under a microscope and counted with a hemocytometer. Cells were then loaded in each channel with a target output of ∼4,000 cells. Reverse transcription and library preparation were performed on C1000 Touch Thermal cycler with 96-Deep Well Reaction Module (Bio-Rad). Amplified cDNA and final libraries were evaluated on a Agilent BioAnalyzer using a High Sensitivity DNA Kit (Agilent Technologies). Individual libraries were diluted to 4nM and pooled for sequencing. Pools were sequenced with 75 cycle run kits (26bp Read1, 8bp Index1 and 55bp Read2) on the NextSeq 500 Sequencing System (Illumina) to ∼70-80% saturation level.

### Pre-processing of scRNA-seq data

ScRNA-Seq data were demultiplexed, aligned to the mouse genome, version mm10, and UMI-collapsed with the Cellranger toolkit (version 2.0.1, 10X Genomics). We excluded cells with fewer than 500 detected genes (where each gene had to have at least one UMI aligned). Gene expression was represented as the fraction of its UMI count with respect to total UMI in the cell and then multiplied by 10,000. We denoted it by TP10K – transcripts per 10K transcripts.

### Dimensionality reduction

We performed dimensionality reduction using gene expression data for a subset of variable genes. The variable genes were selected based on dispersion of binned variance to mean expression ratios using *FindVariableGenes* function of *Seurat* package (110) followed by filtering of cell-cycle, ribosomal protein, and mitochondrial genes. Next, we performed principal component analysis (PCA) and reduced the data to the top 50 PCA components (number of components was chosen based on standard deviations of the principal components – in a plateau region of an “elbow plot”).

### Clustering and sub-clustering

We used graph-based clustering of the PCA reduced data with the Louvain Method (111) after computing a shared nearest neighbor graph (110). We visualized the clusters on a 2D map produced with t-distributed stochastic neighbor embedding (t-SNE) (112). For sub-clustering, we applied the same procedure of finding variable genes, dimensionality reduction, and clustering to the restricted set of data (usually restricted to one initial cluster).

### Differential expression and cluster-specific gene signatures

For each cluster, we used the Wilcoxon Rank-Sum Test to find genes that had significantly different RNA-seq TP10K expression from the remaining clusters (after multiple hypothesis testing correction). As a support measure for ranking differentially expressed genes we also used area under receiver operating characteristic (ROC) curve.

### Filtering out hematopoietic clusters and suspected doublets

Based on cluster annotations with characteristic genes, we removed hematopoietic clusters from further analysis. It is further expected that a small fraction of data should consist of cell doublets (and to an even lesser extent of higher order multiplets) due to co-encapsulation into droplets and/or as occasional pairs of cells that were not dissociated in sample preparation. Therefore, when we found small clusters of cells expressing both hematopoietic and stromal markers we removed them from further analysis. A small number of additional clusters was marked by genes differentially expressed in at least two larger stromal clusters and were annotated as doublets if their average number of expressed genes was higher than the averages for corresponding suspected singlet cluster sources and/or they were not characterized by specific differentially expressed genes. All marked doublets were removed from further analysis.

### Estimation of proliferation status

To score cells for their relative proliferation status, we used a set of characteristic genes involved in cell-cycle (113). For each cell we computed the average expression (TP10K) of cell-cycle genes as a proxy for proliferation status.

### Diffusion maps computation and visualization

We performed non-linear dimensionality reduction of scRNA-seq data by restricting a sparse diffusion matrix of expression data to the eigenspace spanned by eigenvectors corresponding to the top diffusion matrix eigenvalues. We used the *destiny* package (40)), where we used the local estimation of Gaussian kernel width, and the number of nearest neighbors for diffusion matrix approximation was set to the smaller value between the square root of the number of all single cells in the data and 100. The diffusion matrix was constructed on sets of variable genes computed with the same procedure as the one used for clustering and sub-clustering of the data, such that variable genes were re-computed for each diffusion map.

We visualized diffusion maps following the approach described in (32). Specifically, we built a nearest-neighbor graph in the projected space using an implementation of k-NN algorithm from the package *FNN* (114), and then computed a force-directed layout using *ForceAtlas2* (115) from the *Gephi* package (116).

### Connectivity of cell clusters

We quantified the connectivity of single-cell clusters using the partition based graph abstraction method PAGA (117), a part of the single-cell analysis package *Scanpy* (118). The computations were carried out on the same subset of variable genes as for clustering, using default parameters.

### Cell identity assignment in the leukemia scRNA-seq dataset

For scRNA-seq data from control and leukeic mice, we removed hematopoietic cells from the data as described above, and assigned cell types/identities to each cell using differential expression signatures derived from steady state data. Specifically, we collected up to 50 top, most differentially expressed genes in each of the 20 original clusters from the homeostasis data set as signatures. For each candidate cell, we computed its signature scores for ach of the 20 signatures. Each signature score was computed against a background gene set of randomly selected genes. A cell was assigned to the cell-type/cluster with the best signature score. When assigning the sub-cluster identity, we further scored against signature genes of sub-clusters that consisted of up to 20 most differentially expressed sub-cluster genes.

### Changes in cluster sizes in leukemia vs. control

For each cluster, we used the Wald test to quantify the association of condition (control, leukemia) with binomial models. Specifically, for each sample, we collected numbers of cells belonging to a given cluster and the number of cells outside of the cluster. Then we fit a generalized linear model with binomial parameters to the combined data with and without a parameter indicating condition (CTRL, Leukemia). We used an R implementation of Wald test to assess the statistical significance of the difference between the two models. Finally, we corrected the Wald test p-values from all clusters for multiple-hypothesis testing (119).

### Differential expression in leukemia *vs.* control

We observed some contamination of single cell data likely due to ambient RNA. Therefore, prior to computing differentially expressed genes between leukemia and control, we filtered out those genes that were differentially expressed in the hematopoietic contingent of the data when compared with the stromal contingent. The filtered genes were identified separately in leukemia and control conditions, since their respective hematopoietic contingents are known to be different. Specifically, this was achieved by computing differentially expressed genes using Bonferroni corrected Wilcoxon Sum-Rank Test as implemented in *FindAllMarkers* function (default parameters) of the Seurat package and excluding genes from hematopoietic clusters with adjusted p-values < 0.05.

Then, separately for each cluster, we computed genes that were differentially expressed between leukemia and control in two ways. First, we used Bonferroni corrected Wilcoxon Rank-Sum Test (using *FindMarkers* of Seurat package, logfc.threshold=ln(1.2)) to discover differentially expressed genes between conditions. Second, for each cluster, we computed average TP10K expression of cells in every replicate. Then, we used those values in a t-test to assess differences between leukemia and control conditions. This approach mimics bulk RNA-seq measurements. The second approach, although less powerful for discovery of differentially regulated genes, helped us to identify genes that tended to be coherently regulated in samples.

### Gene set enrichment analysis

We used Gene Set Enrichment Analysis (GSEA) with MSigDB (83) gene sets to identify pathways and cellular states having induced or repressed expression in each cell cluster. Specifically, we used the pre-ranked analysis mode, with gene transcripts ranked according to differential expression analysis results (Wilcoxon Runk-Sum Test) of comparing leukemic and control conditions in each cluster. The most significantly over-expressed genes were placed at the top of the ranked list, while the most under-expressed were at the bottom before running the test.

---

**Supplementary figure 1.**
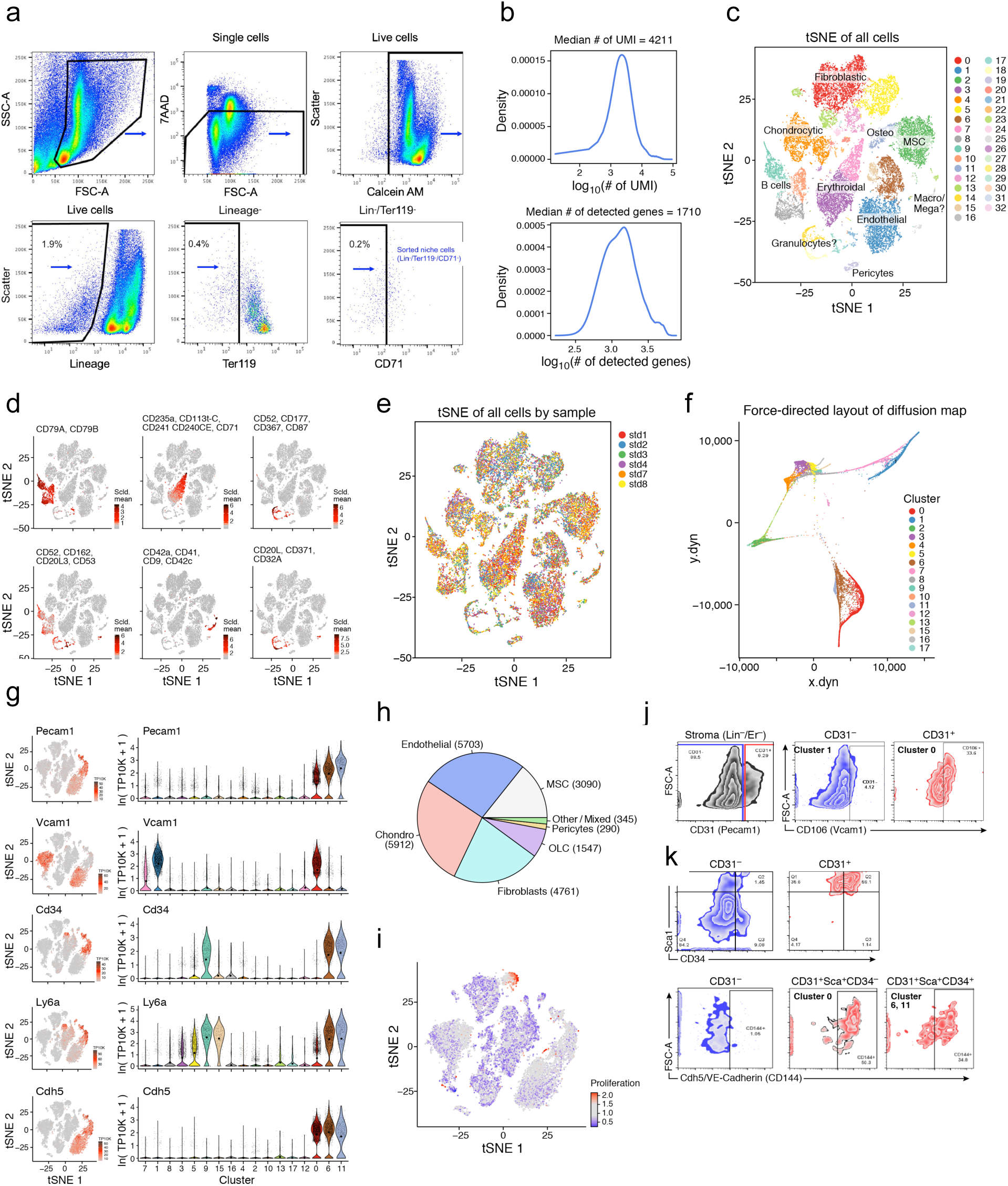
A single cell atlas of the mouse stroma. (**a**) Gating strategy for isolation of mouse bone marrow stroma. (**b**) RNA-seq data quality measures (UMI and genes in cells). (**c**) t-SNE map of the 30,543 bone marrow (BM) cells (hematopoietic and non-hematopoietic) colored by clusters and annotated *post hoc*. (**d**) t-SNE maps as in C) but colored by average expression (color bar, TP10K) of hematopoietic signature genes. (**e**) As in c) but with colors marking samples. (**f**) A force directed layout embedding (FLE) of the cells (dots) from a diffusion map (50 components) computed with the cells from strongly all stroma clusters. (**g**) Signature genes for MSC and ECs. tSNE of **Figure 1b** colored by expression (color bar, TP10K) of key signature genes (left) (right), along with the distributions of expression levels (ln(TP10K+1), *y* axis) for the same genes across the 17 clusters of **Figure 1b** (*x* axis). (**h**) Numbers of BM stromal cells in major cell types. (**i**) tSNE of **Figure 1b** colored by proliferation score (color bar, **Methods**) of select genes used in the field for labeling mesenchymal stem cell populations. (**j**,**k**) FACS analysis of MSCs (cluster 1) and ECs (cluster 0, 6, 11). Same strategy to enrich stroma from immune (Lin-) and erythroid (Er-) cells in (a) was used in combination with antibodies that labels ECs (CD31/*Pecam1*, Sca-1/*Ly6a*, CD34), or MSCs (CD106/*Vcam1*). Cluster 1 cells were separated from all stroma clusters through combination of Lin-/Er-/CD31-/Vcam1+, and cluster 0 through Lin-/Er-/CD31+/Vcam1+. Cluster 6 and 11 were separated from cluster 0 through Lin-/Er-/CD31+/Sca1+/CD34+/VE-Cadherin+.

**Supplementary figure 2.**
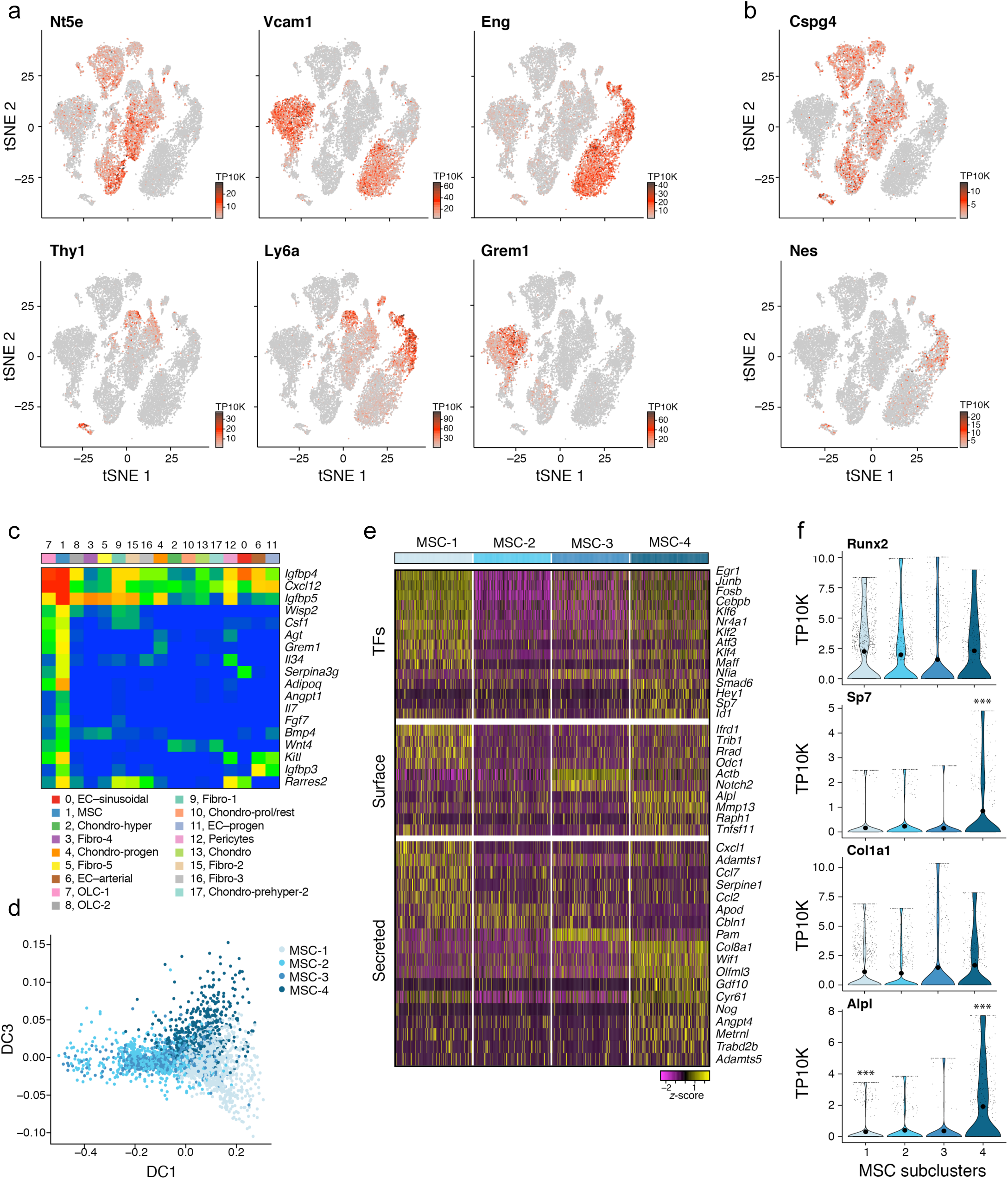
Four subsets of HSC regulator-producing MSCs form a differentiation continuum. (**a**,**b**) tSNE of **Figure 1b** colored by expression (color bar, TP10K) of select genes used in the field for labeling mesenchymal stem cell populations. (**c**) Average expression (ln(TP10K+1)) in all BM stroma clusters of top secreted genes that were also differentially expressed in MSCs (cluster 1). (**d**) MSC sub-clusters diffusion map (2D projection, eigenvectors 1 and 3). (**e**) Top differentially expressed genes among MSC subclusters (single cell view, relative expression z-score). Expression (row-wide z-score of ln of TP10K) of top differentially expressed genes (rows) across the cells (columns) in MSC subclusters. (color bar, top, as in **Figure 2e**), ordered by three gene categories (labels on left). Top differentially expressed genes among MSC sub-clusters (single cell view, relative expression z-score). (**f**) Distributions of expression levels (TP4K, *y* axis, censored scale) for select marker genes across the 4 MSC sub-clusters.

**Supplementary figure 3.**
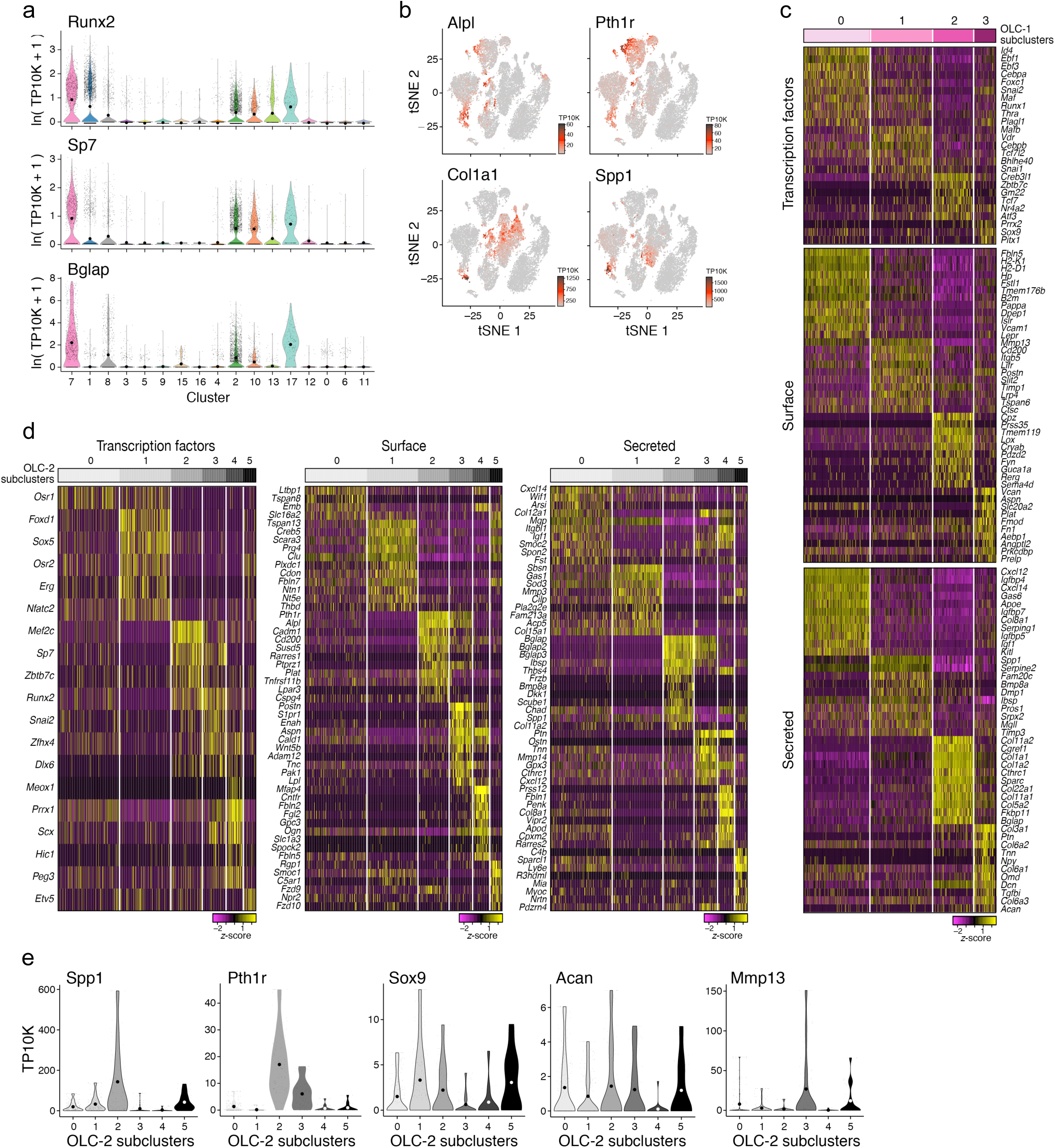
Two osteolineage subsets of distinct differentiation origins and hematopoietic support potential. (**a**) The distributions of expression levels (ln(TP10K+1), *y* axis) for genes as in **Figure 3b** across the 17 clusters of **Figure 1b** (*x* axis). (**b**) tSNE of **Figure 1b** colored by expression (color bar, TP10K) of select OLC related genes. (**c**) Top differentially expressed genes among OLC-1 subclusters (single cell view, relative expression z-score). Expression (row-wide z-score of ln of TP10K) of top differentially expressed genes (rows) across the cells (columns) in OLC-1 subclusters. (color bar, top, as in **Figure 3e**), ordered by three gene categories (labels on left). (**d**) Top differentially expressed genes among OLC-2 subclusters (single cell view, relative expression z-score). Expression (row-wide z-score of ln of TP10K) of top differentially expressed genes (rows) across the cells (columns) in OLC-2 subclusters. (color bar, top, as in **Figure 3k**), ordered by three gene categories (labels on top). (**e**) Distributions of expression levels (TP10K, *y* axis, censored scale) for select marker genes across the 6 OLC-2 sub-clusters.

**Supplementary figure 4.**
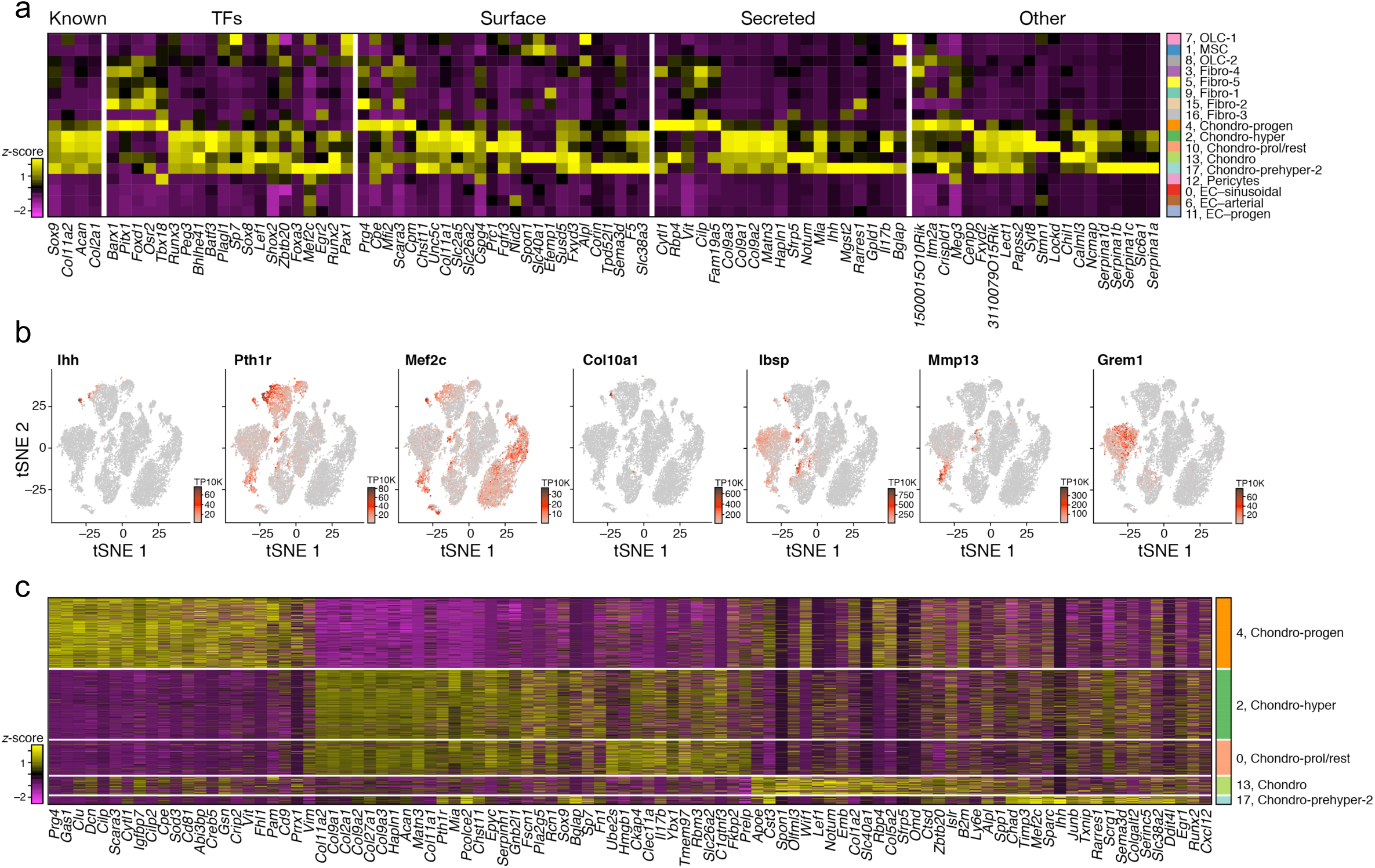
Chondrocyte subsets highlight differentiation paths. (**a**) Expression (column-wide z-score of ln of average TP10K) of top differentially expressed chondrocyte genes (columns) ordered by five gene categories (labels on top) in the cells of each cluster (rows, color bar, right, as in **Figure 1b**). (**b**) tSNE of **Figure 1b** colored by expression (color bar, TP10K) of select genes used for chondrocyte identification. (**c**) Top differentially expressed genes among chondrocyte clusters (single cell view, relative expression z-score). Expression (column-wide z-score of ln of TP10K) of top differentially expressed genes (columns) across the cells (rows) in chondrocyte clusters. (color bar, right, as in **Figure 1d** but only for chondrocyte clusters).

**Supplementary figure 5.**
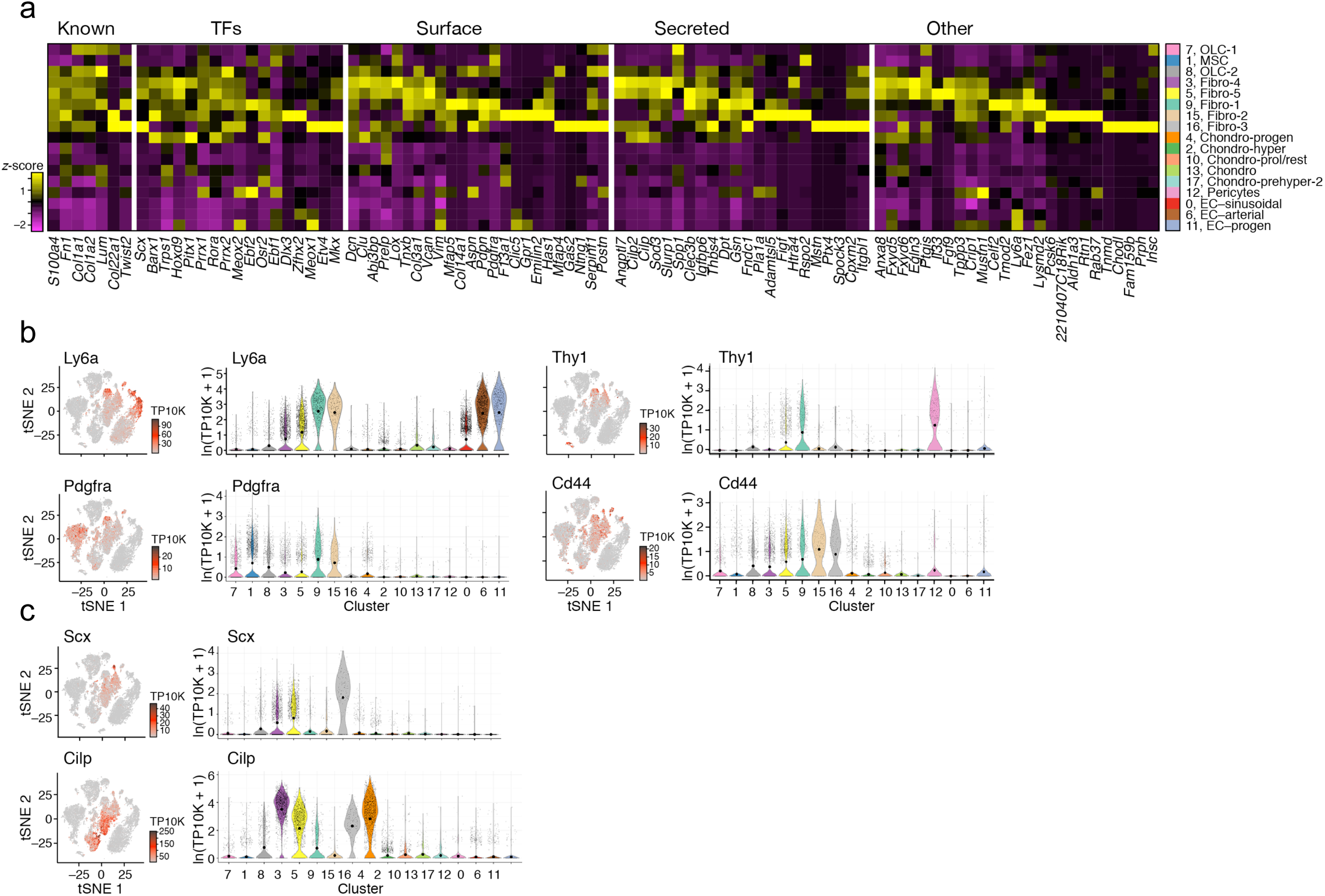
Fibroblasts subsets highlight hematopoiesis support. (**a**) Expression (column-wide z-score of ln of average TP10K) of top differentially expressed fibroblast genes (columns) ordered by five gene categories (labels on top) in the cells of each cluster (rows, color bar, right, as in **Figure 1b**). (**b**,**c**) tSNE of **Figure 1b** colored by expression (color bar, TP10K) of select genes (left) (right), along with the distributions of expression levels (ln(TP10K+1), *y* axis) for the same genes across the 17 clusters of **Figure 1b** (*x* axis).

**Supplementary figure 6.**
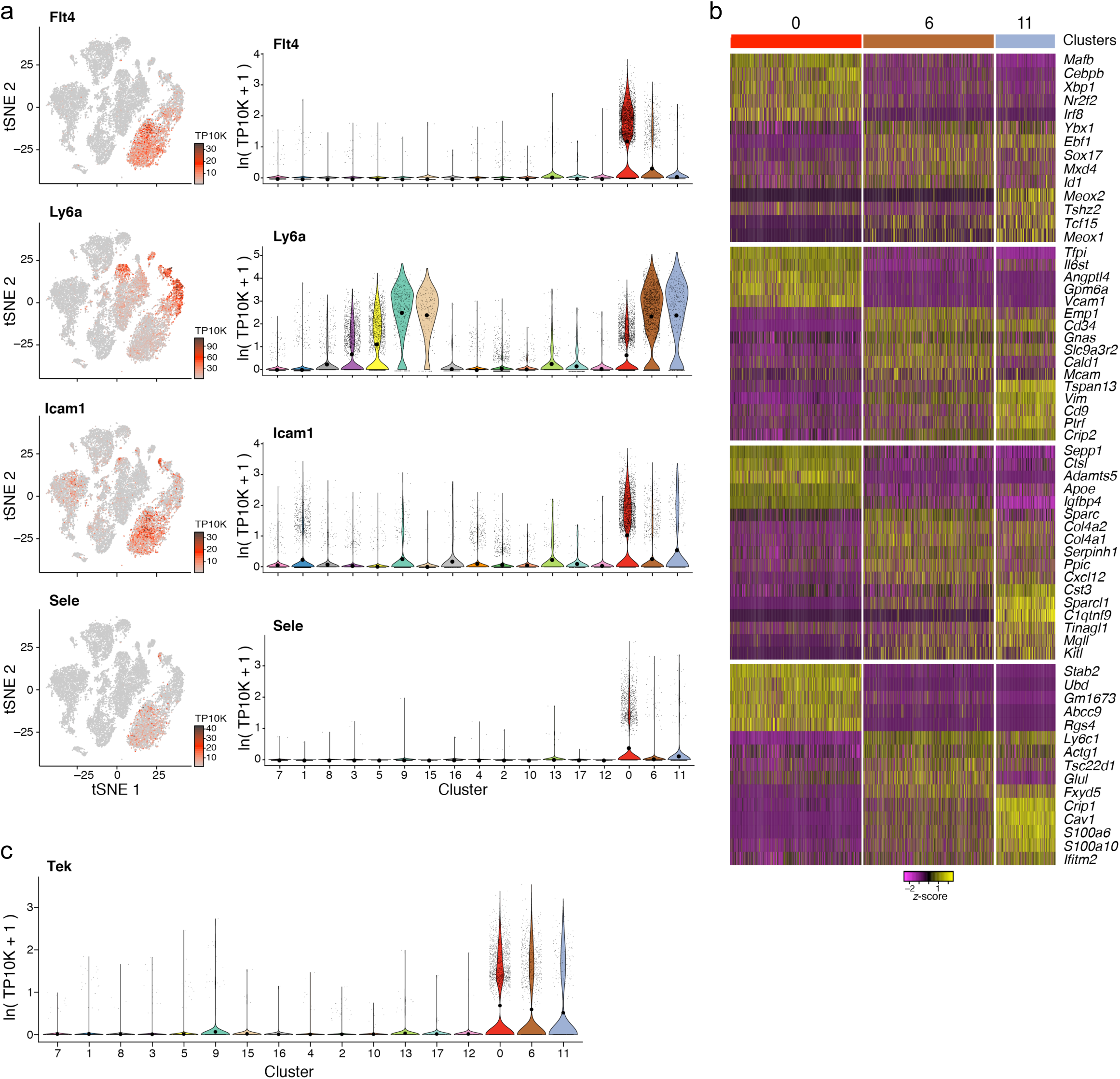
EC progenitors express higher levels of niche factors. (**a**) Signature genes for ECs. tSNE of **Figure 1b** colored by expression (color bar, TP10K) of key EC marker genes (left) (right), along with the distributions of expression levels (ln(TP10K+1), *y* axis) for the same genes across the 17 clusters of **Figure 1b** (*x* axis). (**b**) Top differentially expressed genes among EC clusters (single cell view, relative expression z-score). Expression (row-wide z-score of ln of TP10K) of top differentially expressed genes (rows) across the cells (columns) in EC clusters. (color bar, top, as in **Figure 1d** but only for EC clusters), ordered by four gene categories (labels on left). (**c**) *Tek* gene - the distributions of expression levels (ln(TP10K+1), *y* axis) for the same genes across the 17 clusters of **Figure 1b** (*x* axis).

**Supplementary figure 7.**
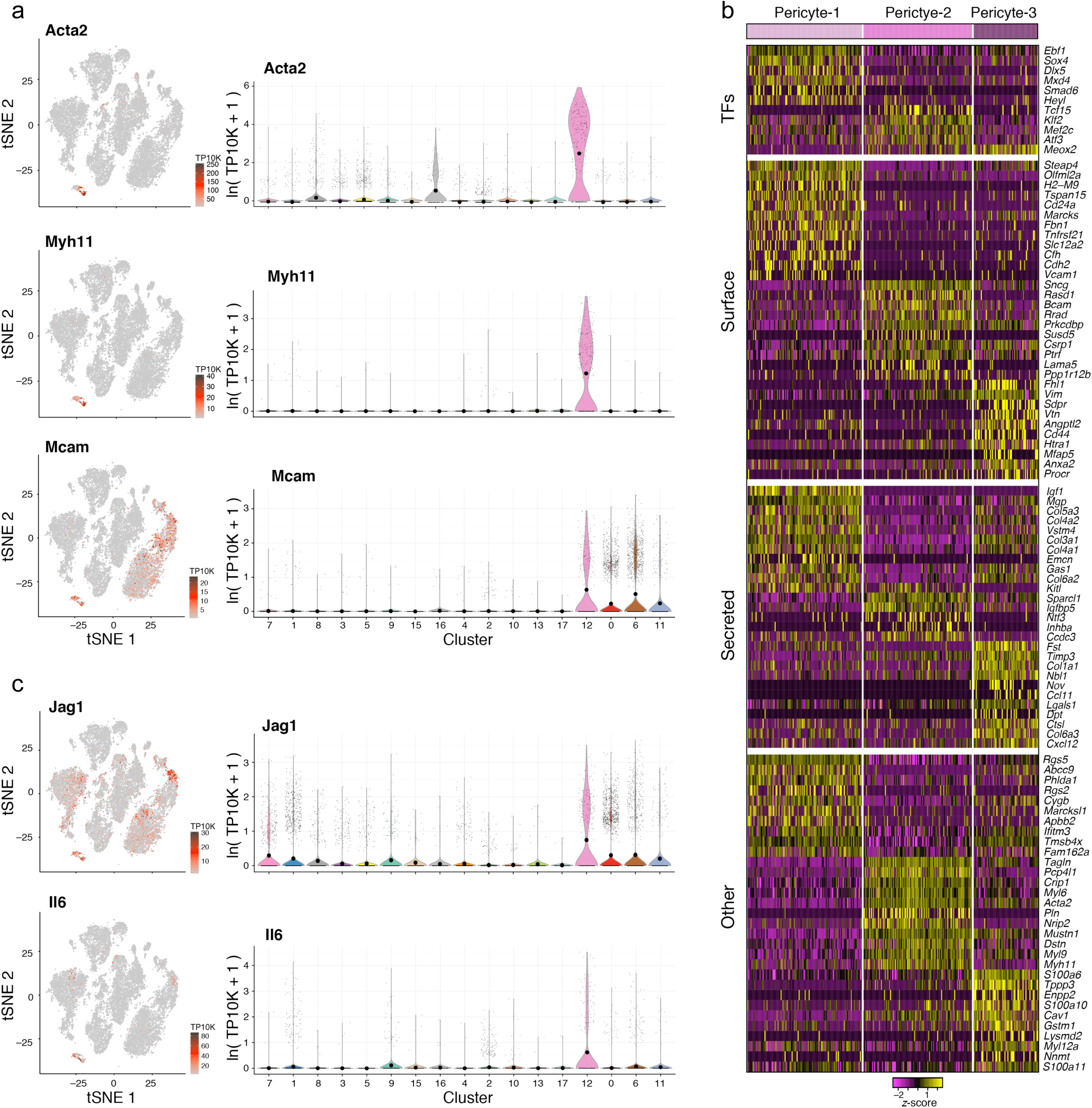
Three distinct subpopulations of pericytes, distinguished from LepR^+^ MSCs. (**a**,**c**) Signature genes for pericytes. tSNE of **Figure 1b** colored by expression (color bar, TP10K) of key marker genes (left) (right), along with the distributions of expression levels (ln(TP10K+1), *y* axis) for the same genes across the 17 clusters of **Figure 1b** (*x* axis). (**b**) Top differentially expressed genes among pericyte sub-clusters (single cell view, relative expression z-score). Expression (row-wide z-score of ln of TP10K) of top differentially expressed genes (rows) across the cells (columns) in pericyte subclusters. (color bar, top, as in **Figure 7d**), ordered by four gene categories (labels on left).

**Supplementary figure 8.**
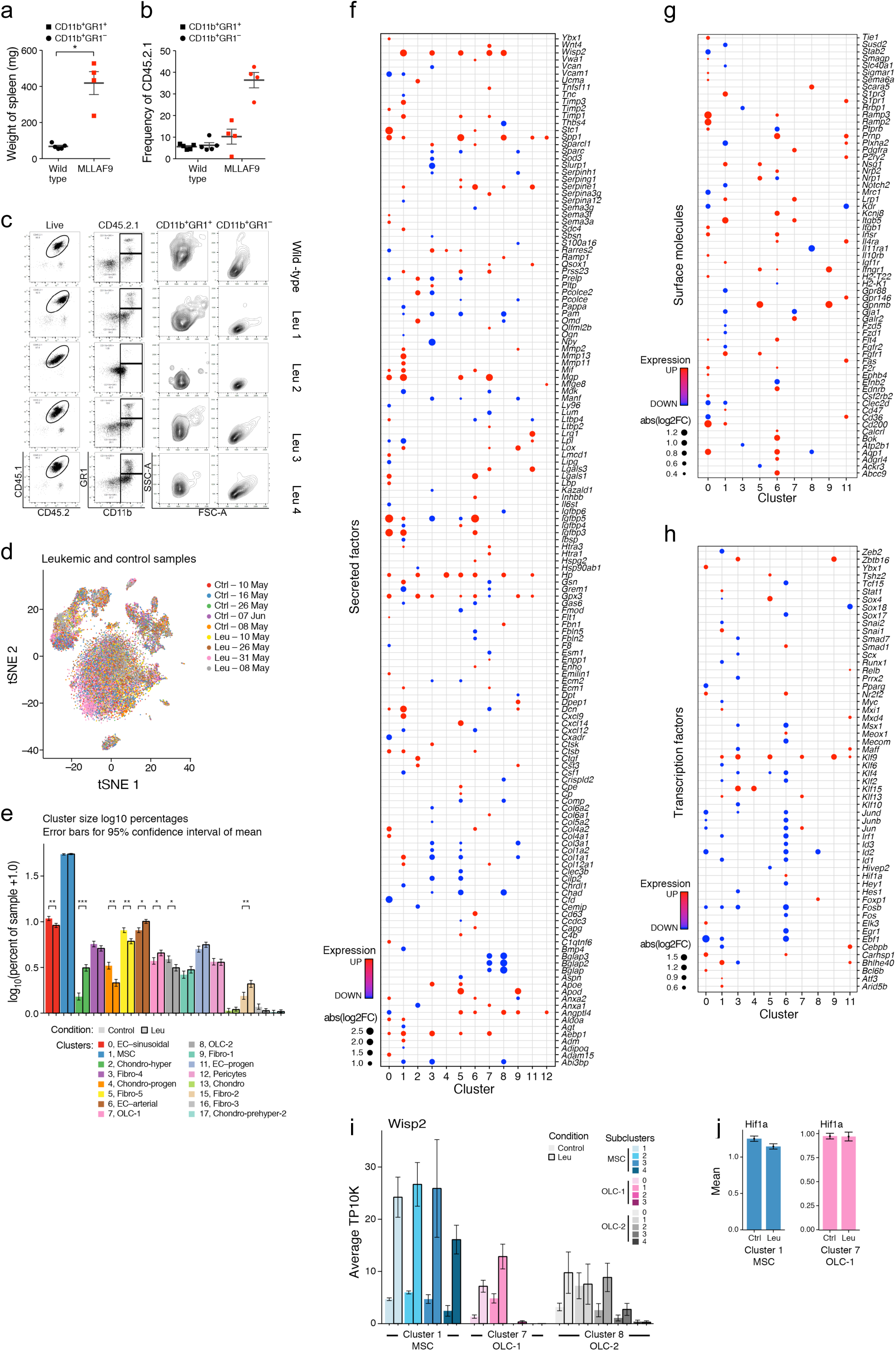
Remodeling of the bone marrow stroma in leukemia. (**a**) Spleen weight of leukemic mice used for single-cell RNA-sequencing experiments. Spleen weights for matched controls (n=5) and leukemic mice (n=4). (student t-test, *p<0.05). (**b**) Donor chimerism and leukemic blast appearance. (**c**) Frequency of myeloid cells in peripheral blood from wild-type control (top row) and MLL-AF9 leukemic mice (row 2-5) characterized by FACS analysis (b). (**d**) A census of the leukemic bone marrow stroma. tSNE of 23,004 cells (dots) from control and leukemic bone marrow colored by sample. (**e**) Compositional changes in BM stroma. Binomial fit mean as percent of cells (y axis) assigned to specific cluster among control or leukemic samples (*x* axis). Error bars: 95% confidence interval of the binomial fit mean. (**f-h**) Fold changes of differentially transcribed genes in leukemia. Adjusted p-value < 0.05), up- (red) and down- (blue) regulated in leukemia. Dot size proportional to base-2 log of fold-change. g) As in F but for cell surface expressed genes. H) As in F but for transcription factor coding genes. (**i**) Changes in *Wisp2* expression in MSCs and OLCs. Average of samples (TP10K, y axis) in MSC and OLC sub-clusters (x axis). Error bars: SEM (**j**) Changes in *Hif1a* expression in MSCs and OLC-1s in leukemia. Average expression (TP10K, y axis). Error bars: SEM.

